# Oxidative Phosphorylation Inhibition in Different Prostate Cancer Models and the Interplay with Androgen Receptor Signaling

**DOI:** 10.1101/2024.09.23.614600

**Authors:** Minas Sakellakis, Sumankalai Ramachandran, Priyanshu Jain, Mark A. Titus

## Abstract

**Introduction:** In prostate cancer (PCa), androgen receptor signaling stimulates both glycolysis and oxidative phosphorylation (OxPhos). Early-stage prostate cancer is particularly reliant on OxPhos for its bioenergetic needs. OxPhos inhibitors have entered clinical trials. Here we investigated their interplay with androgens in different PCa cell lines.

**Methods:** We investigated the effects on PCa cell viability of an ATPase inhibitor (oligomycin) and a complex 1 inhibitor (IACS-010759) in the presence or absence of low testosterone concentrations in vitro. Both androgen-sensitive and insensitive PCa cell lines were used. The effects were assessed using MTT assay, flow cytometry and cell morphology.

**Results:** Treatment with oligomycin resulted in massive apoptotic death of VCAP cells in castrate conditions within 48 hours, but the simultaneous addition of low testosterone levels restored VCAP cell viability. However, complex 1 inhibition with IACS-010759 increased cell viability, which was further promoted in the presence of testosterone. Both oligomycin and IACS-010759 dramatically decreased viability in LNCaP cells, while testosterone had a small but statistically non-significant effect. The antitumor effect of OxPhos inhibitors was smaller in LNCaP-C4-2B compared to LNCaP cells. OxPhos inhibitors slightly decreased proliferation rates in androgen-independent PC3 cells and HEK293 cells. Oligomycin on LNCaP-C4-2B and PC3 cells resulted in an increased number of cells in G0-G1 phase and decreased in S-phase and G2M phase, rather than massive apoptosis.

**Conclusion:** There is an interplay between androgen signaling and OxPhos in androgen-dependent PCa cells. Complex 1 inhibitors should be used with caution, given potential pro-tumorigenic effects in subsets of PCa cells.

## Introduction

Despite recent advances, androgen signaling inhibition (ASI) remains the cornerstone of prostate cancer treatment. In the initial stages of the disease, prostate cancer usually displays a high sensitivity to ASI, in some cases leading to complete disease remission. In such patients, PSA levels may drop to undetectable levels, while a significant proportion of tumor cells may die. However, a small subset of cells typically evade therapy, ultimately leading to disease progression and adverse outcomes [1].

Therefore, targeting and eradicating these residual cells could potentially extend a patient’s life expectancy or even achieve a cure. These cells likely survive anti-androgen treatments either because the therapy does not induce their death but critically stresses them, or because the cells display various degrees of therapy resistance in the first place [2]. Identifying a treatment that synergizes with anti-androgen therapy to enhance antitumor effects could potentially improve patient outcomes dramatically.

Oxidative phosphorylation (OxPhos) occurs inside the mitochondria and is responsible for generating ATP, the bioenergy molecule of cells. The electron transport chain (ETC) is the key player in OxPhos, consisting of five complexes that facilitate redox reactions and create the electrochemical gradient essential for ATP formation. Inhibition of any of these complexes can significantly impact the cellular energy supply, leading to profound consequences [3]. However, the inhibition of individual complexes affects ETC metabolism uniquely. For example, succinate can donate electrons to Complex 2 and bypass the effects of Complex 1 inhibition [4]. Additionally, certain components of Complex 1 play crucial roles in the cellular mechanism of apoptosis [5].

Cancer cells that survive OxPhos inhibition usually compensate for the loss of bioenergy (ATP) production by up-regulating glycolysis. In prostate cancer, androgens stimulate both glycolysis and OxPhos by activating androgen receptor (AR) signaling [6]. Interestingly, early-stage prostate cancer has been found to rely more on OxPhos for its bioenergetic needs, while more advanced disease shifts towards increased reliance on glycolysis, resembling other aggressive tumors exhibiting the Warburg effect [7]. Here we investigated the heterogeneity of OxPhos inhibition in different androgen-responsive and non-responsive prostate cancer cell lines in vitro. We also hypothesized that androgen signaling might act as a mechanism that facilitates the survival of androgen-dependent prostate cancer cells

exposed to OxPhos inhibition. Androgen deprivation and OxPhos inhibition could potentially synergistically enhance antitumor effects in specific subsets of prostate cancer tumor cells. We also hypothesized that different levels of OxPhos inhibition will have different effects in various prostate cancer models.

## Results

### Effects on VCaP cells

The proliferation rate of VCAP cells (which are known to harbor amplified AR) in charcoal stripped serum (CSS) media versus control conditions (fetal bovine serum media) at 72 hours was assessed. As expected, CSS conditions negatively impacted VCAP cell viability (p<0.001)(Fig. 1A). The treatment with 50 to 100 ng/mL (0.06-0.12 nM) of oligomycin (O) caused nearly complete eradication of VCAP cells in CSS within 48 hours (p<0.001 for both conditions versus CSS alone). However, the simultaneous addition of low testosterone (T) levels (80 ng/dL or 2.8 nM) rescued the cells, leading to a restoration of VCAP cell viability to levels higher than those observed with CSS alone (p<0.001 against CSS alone, p=0.5 against CSS with T)(Fig. 1A). These results strongly suggest that AR signaling inhibition is a critical mediator of oligomycin-induced toxicity in VCAP cells. Concurrent AR signaling inhibition and oxidative phosphorylation (OxPhos) inhibition was lethal to VCAP cells. Moreover, the addition of oligomycin in CSS significantly increased apoptotic rates in VCAP cells compared to the other conditions (p=0.021 against CSS and T, p=0.027 against CSS and T and O), further supporting the synergistic anti-tumor effect (Fig. 1B and 1C). We did not observe other major cell cycle effects with use of oligomycin.

**Figure 1A.** Growth of VCAP cells after 72 hours under the absence or presence of castration (FBS vs CSS media), low testosterone levels (80 ng/dL or2.77 nM), 50-100 ng/ml (0.06-0.12 nM) oligomycin, or combinations thereof. Lines represent mean values and standard deviations, while dots represent light absorbance measurement at 570 nm (MTT assay). FBS=fetal bovine serum, CSS=charcoal-stripped serum

**Figure 1B.** Flow cytometry results depicting percentage of cells in Sub-G1 phase (a) and cell cycle changes (b) in VCAP after 48 hours in the absence or presence of castration (FBS or CS media), presence or absence of 80 ng/dL (or 2.77nM) testosterone, presence or absence of 50 ng/ml (0.06 nM) oligomycin, or their combinations. FBS=fetal bovine serum, CS=charcoal-stripped serum, O=oligomycin, T=testosterone

**Figure 1C.** Morphological changes in VCaP cells after 48 hours in the presence of castration (CSS), 80 ng/dL (2.77 nM) testosterone and 50 ng/ml (0.06 nM) oligomycin, or their combinations. CSS=charcoal- stripped serum, O=oligomycin, T=testosterone

**Figure 1D.** Growth of VCAP cells after 72 hours under the absence or presence of castration (FBS vs CSS media), low testosterone levels (80 ng/dl or 2.77nM), 25 nM IACS-010759, or combinations thereof. Lines represent mean values and standard deviations, while dots represent light absorbance measurement at 570 nm (MTT assay). FBS=fetal bovine serum, CSS=charcoal-stripped serum, nM=nanomolar

### Effects on LNCaP cells

Next, we performed the experiments using LNCaP cells, which are androgen-sensitive without AR amplification. As expected, CSS conditions negatively impacted LNCaP cell viability compared to control. Once again, the addition of 0.06 nM oligomycin caused almost complete elimination of LNCaP cells within 72 hours in CSS media, as was demonstrated by MTT and morphological studies (p<0.001)(Fig. 2A). When 2.77 nM testosterone was concurrently added, a small not statistically significant increase in LNCaP cell viability was observed (p=0.079), and cell viability was well below that of CSS alone (p<0.001)(Fig. 2A). These data suggest that LNCaP cells are highly sensitive to ATP synthase inhibition even in the presence of testosterone. The addition of oligomycin increased apoptosis in CSS treated and slightly increased apoptosis in CSS plus testosterone-treated LNCaP cells (p=0.004 and p=0.05 respectively). Morphological studies suggest non-apoptotic cell death plays a role as well. We also observed a substantial percentage of LNCaP cells in G0-G1 phase (84-87%) and decreased S-phase and G2M phase (p=0.006, p=0.019 and 0.048 respectively)(Fig. 2B and C). This effect was observed even in the presence of testosterone (p=0.01, 0.019 and p=0.006 respectively).

**Figure 2A.** Growth of LNCaP cells after 72 hours under the absence or presence of castration (FBS vs CSS media), low testosterone levels (80 ng/dL or 2.77 nM), 50-100 ng/ml (0.06-0.12 nM) oligomycin, or combinations thereof. Lines represent mean values and standard deviations, while dots represent light absorbance measurement at 570 nm (MTT assay). FBS=fetal bovine serum, CSS=charcoal-stripped serum

**Figure 2B.** Flow cytometry results depicting the percentage of cells in Sub-G1 phase (a) and cell cycle changes (b) in LNCaP after 48 hours in the absence or presence of castration (FBS or CS media), presence or absence of 80 ng/dL (2.77 nM) testosterone, presence or absence of 50 ng/ml (0.06 nM) oligomycin, or their combinations. FBS=fetal bovine serum, CS=charcoal-stripped serum, O=oligomycin, T=testosterone.

**Figure 2C.** Morphological changes in LNCaP cells after 48 hours in the presence of castration (CSS), 80 ng/dL (2.77 nM) testosterone and 50 ng/ml (0.06 nM) oligomycin, or their combinations. CSS=charcoal- stripped serum, O=oligomycin, T=testosterone.

**Figure 2D.** Growth of LNCaP cells after 72 hours under the absence or presence of castration (FBS vs CSS media), low testosterone levels (80 ng/dL or 2.77 nM), 25 nM IACS-010759, or combinations thereof. Lines represent mean values and standard deviations, while dots represent light absorbance measurement at 570 nm (MTT assay). FBS=fetal bovine serum, CSS=charcoal-stripped serum, nM=nanomolar

The inhibition of Complex 1 with 25 nM IACS-010759 under CSS conditions also dramatically reduced LNCaP cell viability (p<0.001). The concurrent addition of small amounts of testosterone in the presence of IACS-010759 was unable to rescue the tumor cells and even caused a further marginal decrease in viability (p=0.045)(Fig. 2E1 and 2).

### Effects on LNCaP-C4-2B cells

LNCaP-C4-2B cells are an isogenic cell line to LNCaP, characterized by persistent AR-induced transcription when grown in CSS conditions [8]. LNCaP-C4-2B cell viability in CSS media decreased compared to control. The addition of 0.06 nM oligomycin in CSS decreased LNCaP-C4-2B cell viability, although not to the extent observed in LNCaP cells, as the cells continued to proliferate after 72 hours (p<0.001). Concurrent addition of low testosterone levels resulted in a small but significant restoration of LNCaP-C4-2B cell viability, albeit less compared to cells in CSS alone (p<0.001)(Fig. 3A). These findings suggest that the persistent AR-mediated transcription in LNCaP-C4-2B cells might have a small protective effect against oligomycin toxicity.

**Figure 3A.** Growth of LNCaP-C4-2B cells after 72 hours under the absence or presence of castration (FBS vs CSS media), low testosterone levels (80 ng/dL or 2.77 nM), 50-100 ng/ml (0.06-0.12 nM) oligomycin, or combinations thereof. Lines represent mean values and standard deviations, while dots represent light absorbance measurement at 570 nm (MTT assay). FBS=fetal bovine serum, CSS=charcoal-stripped serum

During oligomycin treatment, overall cellular viability after 72 hours was higher compared to the beginning of the experiment, which could be attributed mostly to a decrease in proliferation and a slower growth rate, rather than the substantial cellular apoptosis observed in VCAP cells (Fig. 3A). Cell cycle changes were similar to LNCaP cells with increased G0-G1 (p=0.008) and decreased S (p=0.025) and G2M phase (p=0.167), although a smaller absolute percentage of cells were in G0-G1 phase and higher in G2M phase compared to LNCaP (Fig. 3B). This can also be evidenced by studying the morphological changes of LNCaP-C4-2B cells under the microscope (Fig. 3C).

**Figure 3B.** Flow cytometry results depicting the percentage of cells in Sub-G1 phase (a) and cell cycle changes (b) in LNCaP-C4-2B after 48 hours in the absence or presence of castration (FBS or CS media), presence or absence of 80 ng/dL (2.77 nM) testosterone, presence or absence of 50 ng/ml (0.06 nM) oligomycin, or their combinations. FBS=fetal bovine serum, CS=charcoal-stripped serum, O=oligomycin, T=testosterone

**Figure 3C.** Morphological changes in LNCaP-C4-2B cells after 48 hours in the presence of castration (CSS), 80 ng/dL (2.77 nM) testosterone and 50 ng/ml (0.06 nM) oligomycin, or their combinations. CSS=charcoal-stripped serum, O=oligomycin, T=testosterone

**Figure 3D.** Growth of LNCaP-C4-2B cells after 72 hours under the absence or presence of castration (FBS vs CSS media), low testosterone levels (80 ng/dL or 2.77 nM), 25 nM IACS-010759, or combinations thereof. Lines represent mean values and standard deviations, while dots represent light absorbance measurement at 570 nm (MTT assay). FBS=fetal bovine serum, CSS=charcoal-stripped serum, nM=nanomolar

Paradoxically, the addition of 25 nM IACS-010759 under CSS media conditions dramatically decreased LNCaP-C4-2B cell viability (p<0.001)(Fig. 3E). Concurrent addition of 2.77 nM testosterone failed to salvage the cells (p=0.076).

### Effects on PC3 cells

PC3 cells represent advanced metastatic disease with AR-insensitive characteristics. Treatment with 0.06 nM oligomycin minimally decreased PC3 cell proliferation (p<0.001). PC3 cells were the least affected PCa cell lines treated with oligomycin. As expected, due to their AR-insensitive status, the concurrent addition of low testosterone levels had no effect on this rapidly proliferating cell line (p=0.654)(Fig. 4A). The negative impact of oligomycin after 72 hours was attributed mostly to an overall decrease in proliferation and slowing growth rate, rather than massive cellular death. Oligomycin’s impact mainly occurred through cell cycle alterations other than apoptosis induction. Specifically, oligomycin increased the percentage of cells in G0-G1 phase and decreased the percentage of cells in S phase and G2M phase (Fig. 4B). These conclusions were further supported by the morphological features of the treated cells (Fig. 4C). Similar to oligomycin, Complex 1 inhibition with 25 nM IACS-010759 resulted in a small decrease in PC3 cell viability and proliferation (p<0.001), while the concurrent addition of small testosterone levels did not restore cell viability, as expected (p=0.826)(Fig. 4E).

**Figure 4A.** Growth of PC3 cells after 72 hours under the absence or presence of castration (FBS vs CSS media), low testosterone levels (80 ng/dL or 2.77 nM), 50-100 ng/ml (0.06-0.12 nM) oligomycin, or combinations thereof. Lines represent mean values and standard deviations, while dots represent light absorbance measurement at 570 nm (MTT assay). FBS=fetal bovine serum, CSS=charcoal-stripped serum

**Figure 4B.** Flow cytometry results depicting the percentage of cells in Sub-G1 phase (a) and cell cycle changes (b) in PC3 after 48 hours in the absence or presence of castration (FBS or CS media), presence or absence of 80 ng/dL (2.77 nM) testosterone, presence or absence of 50 ng/ml (0.06 nM) oligomycin, or their combinations. FBS=fetal bovine serum, CS=charcoal-stripped serum, O=oligomycin, T=testosterone

**Figure 4C.** Morphological changes in PC3 cells after 48 hours in the presence of castration (CSS), 80 ng/dL (2.77 nM) testosterone and 50 ng/ml (0.06 nM) oligomycin, or their combinations. CSS=charcoal- stripped serum, O=oligomycin, T=testosterone

**Figure 4D.** Growth of PC3 cells after 72 hours under the absence or presence of castration (FBS vs CSS media), low testosterone levels (80 ng/dl or 2.77 nM), 25 nM IACS-010759, or combinations thereof. Lines represent mean values and standard deviations, while dots represent light absorbance measurement at 570 nm (MTT assay). FBS=fetal bovine serum, CSS=charcoal-stripped serum, nM=nanomolar

### Effects on HEK293 cells

HEK293 cells are normal human embryonic kidney cells and were used as a non-cancer AR negative and androgen-indifferent control model. The addition of 0.06 nM oligomycin under CSS conditions decreased HEK293 cell viability (p-value<0.001), but viability after 72 hours was increased compared to cells at 0 hours (p<0.001). The concurrent addition of 2.77 nM testosterone had no effect on cell viability (p=0.957)(Fig. 5A). Oligomycin’s impact on cell viability and proliferation occurred mostly through cell cycle alterations other than apoptosis (p=0.124 for apoptosis)(Fig. 5B). Combined with morphological features, these data suggest that the effect of oligomycin is likely mostly attributed to decreased proliferation rather than massive cell death (p=0.079 for G0/G1, p=0.064 for S, p=0.226 for G2/M)(Fig. 5C). Similar to oligomycin, the addition of 25 nM IACS-010759 in all conditions decreased HEK293 cell viability (p<0.001), but not to the extent of LNCaP or LNCaP-C4-2B cells, while the concurrent addition of testosterone had no effect, as anticipated (p=0.852)(Fig. 5E).

**Figure 5A.** Growth of HEK293 cells after 72 hours under the absence or presence of castration (FBS vs CSS media), low testosterone levels (80 ng/dL or 2.77 nM), 50-100 ng/ml (0.06-0.12 nM) oligomycin, or combinations thereof. Lines represent mean values and standard deviations, while dots represent light absorbance measurement at 570 nm (MTT assay). FBS=fetal bovine serum, CSS=charcoal-stripped serum

**Figure 5B.** Flow cytometry results depicting the percentage of cells in Sub-G1 phase (a) and cell cycle changes (b) in HEK293 after 48 hours in the absence or presence of castration (FBS or CS media), presence or absence of 80 ng/dL (2.77 nM) testosterone, presence or absence of 50 ng/ml (0.06 nM) oligomycin, or their combinations. FBS=fetal bovine serum, CS=charcoal-stripped serum, O=oligomycin, T=testosterone

**Figure 5C.** Morphological changes in LNCaP cells after 48 hours in the presence of castration (CSS), 80 ng/dL (2.77 nM) testosterone and 50 ng/ml (0.06 nM) oligomycin, or their combinations. CSS=charcoal- stripped serum, O=oligomycin, T=testosterone

**Figure 5D.** Growth of HEK293 cells after 72 hours under the absence or presence of castration (FBS vs CSS media), low testosterone levels (80 ng/dL or 2.77 nM), 25 nM IACS-010759, or combinations thereof. Lines represent mean values and standard deviations, while dots represent light absorbance measurement at 570 nm (MTT assay). FBS=fetal bovine serum, CSS=charcoal-stripped serum, nM=nanomolar

### Tricarboxylic acid (TCA) metabolite levels

We developed a platform to study the TCA metabolite levels after OxPhos inhibition in castrate and non-castrate environments in vitro by using liquid chromatography mass spectrometry.

The presence of oligomycin in VCaP cells tended to decrease the amount of detected pyruvate (Supplementary file). Moreover, no increased succinate amounts were observed. Interestingly, the addition of IACS-010759 tended to increase the measured lactate amounts. In LNCaP cells no consistent metabolite shifts were measured after addition of oligomycin. It is unknown if direct tumor death by necrosis plays a role. Given the toxicity of oligomycin in LNCaP cells, this suggests a lack of metabolic shift or metabolic adaptation. However, the wide range of results should warrant caution during interpretation. Lactate concentrations tended to be low after the addition of IACS-010759 in LNCaP cells. Similar patterns were observed in LNCaP-C4-2B cells. No dramatic shifts in metabolites were also observed after the addition of oligomycin in PC3 cells, which were the cell lines that had the highest proliferation rates and were the least affected by oligomycin. Unfortunately, this metabolite platform is a work-in-progress and the results are preliminary and not definite. Several challenges remain, including how to deal with the presence of both dead and alive cells in the collected pellets, protocol optimization, the complexity of adjusting parameters based on different cell counts etc.

## Discussion

Since the discovery of the Warburg effect, extensive research efforts have been directed towards inhibiting glycolysis [7]. However, accumulating evidence suggests that certain tumors can also heavily rely on OxPhos for survival. Increased dependency on OxPhos is often a hallmark of cancer cells with stem cell properties, specific mutations, or resistance to targeted therapy and chemotherapy [10, 11]. Consequently, OxPhos has emerged as an attractive target for novel antitumor agents.

Targeting OxPhos has proven to be challenging, as many investigational drugs have shown increased toxicity without a therapeutic window, since many healthy cells also depend on OxPhos for survival [12]. Therefore, the ideal drug candidate should display high potency and increased selectivity against tumor cells. For example, a few mitochondrial uncouplers exhibit increased cytotoxicity against cancer cells due to mitochondrial hyperpolarization or a more alkaline mitochondrial matrix in these cells [13].

Another strategy involves using agents that selectively render cancer cells highly sensitive to OxPhos inhibitors, with which they would act synergistically. In this scenario, lower doses of a potentially toxic drug would be active against vulnerable or stressed tumor cells, while sparing normal cells from significant effects.

Several complex 1 inhibitors have entered clinical trials, but the results so far have been underwhelming. In a recent phase 1 clinical study, IACS-010759 showed activity in 2 patients with prostate cancer, but overall, the drug demonstrated low selectivity against tumor cells and increased neurotoxicity [12, 14]. Furthermore, our study revealed that the effect on a subset of prostate cancer cells might be contrary to what was anticipated. While IACS-010759 was highly toxic for LNCaP and LNCaP-C4-2B cells, it promoted tumor cell proliferation in VCaP cells, even in the absence of testosterone. This effect was not observed with oligomycin, suggesting that some Complex 1 inhibitors may even be detrimental when used against specific prostate cancer subtypes, especially without concurrent androgen deprivation. Moreover, they can potentially result in mixed effects against different subclones in the case of highly heterogeneous tumors.

In this study, we showed that androgen deprivation acts synergistically with OxPhos inhibition in specific subsets of prostate cancer cells. Notably, although each treatment alone had a modest effect, the combination of androgen deprivation and ATP synthase inhibition caused massive apoptosis in VCaP cells. These findings suggest an intricate interplay between OxPhos activity and androgen signaling in VCaP cells. Caution should be exercised when employing Complex 1 inhibitors against cells that share properties similar to VCaP cells. However, it is worth noting that VCaP cells were unable to survive long-term in CSS alone, even in the presence of IACS-010759.

On the other hand, LNCaP cells exhibited extremely high sensitivity to OxPhos inhibition, while the addition of testosterone had a small pro-survival effect (albeit not statistically significant). LNCaP-C4-2B

cells in general displayed less sensitivity than LNCaP cells to ATP synthase inhibition, suggesting that the presence of androgen signaling in the absence of androgens in these cells might confer some degree of resistance to OXPHOS inhibition. However, LNCaP-C4-2B were extremely sensitive to 25 nM IACS- 010759, even in the presence of testosterone.

The antitumor effect of oligomycin in LNCaP-C4-2B and PC3 cells was mostly achieved from an increased number of cells remaining in G0-G1 phase and decreased in S-phase and G2M phase, which resulted in decreased proliferation rather than massive death. This implicates a differential effect of OxPhos inhibition depending on the androgen-sensitivity status, although this may vary according to specific enzyme of the mitochondrial respiratory chain that is inhibited.

The investigation of concentration changes in TCA metabolites during blockage of different respiratory chain complexes holds promise for providing further mechanistic explanations. Moreover, it can suggest ways to overcome treatment failures of Complex 1 and other inhibitors. For example, the addition of testosterone synergistically enhanced proliferation of VCaP cells when Complex 1 was inhibited.

Interestingly, this was associated with increased lactate production. It is well known that increased lactate can pose a pro-tumorigenic effect by entering the tumor cells and sustaining mitochondrial metabolism [16]. Complex 1 inhibition can be bypassed by increased production of TCA metabolites such as succinate, which can be theoretically achieved when increased amounts of lactate are converted to pyruvate and enter the TCA cycle. Notably, our platform on metabolite concentrations is a work in-progress and the findings should be interpreted with caution.

Limitations of our study include the reliance solely on in vitro evidence, the short term follow up in the experiments, and the use of only two OxPhos inhibitors. Although mitochondrial inhibition may interfere with the interpretation of MTT data, the results were not only clear but were also validated with other methods, for example with morphological studies. Despite the limitations, our study provides a strong foundation to answer critical questions regarding the use of OxPhos inhibitors in prostate cancer. The latter have already entered clinical trials, although extensive pre-clinical knowledge regarding their activity and interactions with other drugs or pathways is still lacking.

## Conclusions

In conclusion, our findings highlight the interplay between androgen signaling and OxPhos in androgen-dependent prostate cancer cells. It is crucial to exercise caution and avoid the use of OxPhos inhibitors (especially complex I inhibitors) as standalone treatments against prostate cancer. A more reasonable therapeutic approach involves the dual inhibition of androgen signaling and OxPhos, which warrants further investigation.

## Materials and methods

### Materials

LNCaP, HEK293 and PC3 cells were obtained from American Type culture Collection (ATCC). LNCaP-C4- 2B and VCaP cells were kind gifts from Dr. Nora Navone and Dr. Timothy Thompson, respectively.

Roswell Park Memorial Institute (RPMI) 1640 Medium (Cat# SH30027.01), Dulbecco’s modified Eagle’s medium (DMEM, Cat# SH30243.01) and DPBS (Cat# SH30378.02) were purchased from Hyclone (San Angelo, TX). Fetal bovine serum (FBS, Cat# 900-108) and charcoal:dextran stripped FBS (CSS, Cat# 100.119) were purchased from Gemini Bioproducts (Sacramento, CA). Thiazolyl blue tetrazolium bromide (Cat# M5655), DMF (Cat# 4551), sodium dodecyl sulfate (SDS)(Cat#slbt3991), acetic acid (Cat# 695092) and trypsin (Cat# 59417C) were obtained from Sigma-Aldrich (St Louis, MO). Testosterone (Cat# S6066SDS) was purchased from Isosciences (Ambler, PA). Penicillin/Streptomycin (Cat# 30-002-CI) and Hepes buffer (Cat# 25-060-CI) were purchased from Corning (NY, USA). N-(3-Dimethylaminopropyl)-N-ethylcarbodiimide was purchased from sigma-aldrich (Cat#BCBL6239V). UHPLC-MS W8-1 Water (Cat#182518) and Methanol (Cat#172940) were purchased from thermo fisher scientific (Belgium, EU). Pyridine (Cat#434871/1) was purchased from Fluka Chemika (Buchs, Switzerland, EU). 3- Nitrophenylhydrazine hydrochloride (Cat#S54248V) was purchased from sigma-aldrich (St. Louis, MO). Propidium Iodine (Cat#1932759) was purchased from Invitrogen (Carlsbad, CA, USA). Oligomycin (Cat#SLBS6501) was purchased from sigma-aldrich (St. Louis, MO). RNAse (Cat#85123821) was purchased from Boehringer Mannheim (Germany, EU).

### Cell growth and viability assay

Experiments were conducted in 5 prostate cancer cell lines, VCaP, LNCaP, LNCaP-C4-2B, PC3, and HEK293 cells. Equal numbers of cells were seeded initially in a 96-well plate, each well containing 100uL media. The cells were grown for 24 hours in Roswell Park Memorial Institute (RPMI) 1640 Medium with 10% Fetal Bovine Serum (FBS) and 1% penicillin/streptomycin (PS) media. For VCaP and HEK293 cells we used Dulbecco Modified Eagle Medium (DMEM) instead of RPMI 1640. After 24 hours the old media was replaced with charcoal-stripped serum -containing (CSS) media alone or with the addition of 2.7 nM testosterone and the OXPHOS inhibitors oligomycin and IACS-010759. For IACS-010759 we used a clinically relevant dose of 5nM. Given the lack of clinical use of oligomycin, we used 50-100 ng/ml, a concentration that is able to disrupt the activity of ATP synthase. After 72 hrs we performed 3-(4,5- dimethylthiazol-2-yl)-2,5-diphenyl-2H-tetrazolium bromide (MTT) assays using the Roche MTT protocol (Sigma Aldrich Cat#11465007001). After 24 hours light absorption at 570 nm was measured using the MTT plate reader (Wallac EnVision 1.12). Independent sample t-tests were used to assess for differences in light absorption. All experiments were performed in triplicate.

### Fluorescence-Activated Cell Sorting

VCaP and HEK cells were initially seeded in DMEM/FBS/penicillin-streptomycin media. LNCaP, LNCaP-C4- 2B and PC3 cells were seeded in RPMI 1640/FBS/penicillin-streptomycin media. After 24 hours the media was replaced with new media containing the studied conditions. After 48 hours, 2 million cells were fixed and stained for propidium iodine. Cell cycle distribution was analyzed with a FACS Calibur (Becton Dickinson, Bedford, MA).

### Cell Morphology

VCaP and HEK cells were seeded in DMEM/FBS/penicillin-streptomycin media and LNCaP, LNCaP-C4-2B and PC3 cells were seeded in RPMI 1640/FBS/penicillin-streptomycin media for 24 hours. The media was then replaced with new media that contained studied conditions. After 48 hours, images were captured using an Olympus CK-40-F100 microscope.

### Mass Spectrometry

After an equal number of cells were placed for 48 hours in the studied conditions media, cell pellets were collected and analyzed for pyruvate and lactate concentrations, using liquid chromatography-tandem mass spectrometry (LCMS) as previously described, and normalized for cell count [6].

### Statistical analysis

Independent sample t-tests were used for comparisons of different conditions. All statistical analyses were performed using SPSS 26 version.

### Clinical trial registration

This study does not report the results of a clinical trial.

### Patient consent

This study did not report clinical data, hence patient consent was not needed.

## Author contributions

Concept and design: Minas Sakellakis and Mark A. Titus.

Collection and assembly of data: Minas J Sakellakis, Sumankalai Ramachandran, Priyanshu Jain and Mark A. Titus.

Data analysis and interpretation: All authors. Manuscript writing: All authors.

Final approval of manuscript: All authors.

## Ethics approval

The authors obtained approval by the University of Texas MD Anderson Cancer Center Institutional Review Board.

## Acknowledgement

No acknowledgements

## Data availability statement

Data is available upon request to the corresponding author.

## Supplementary file

The effect of OxPhos inhibition on pyruvate, lactate, succinate, malate and fumarate levels

---

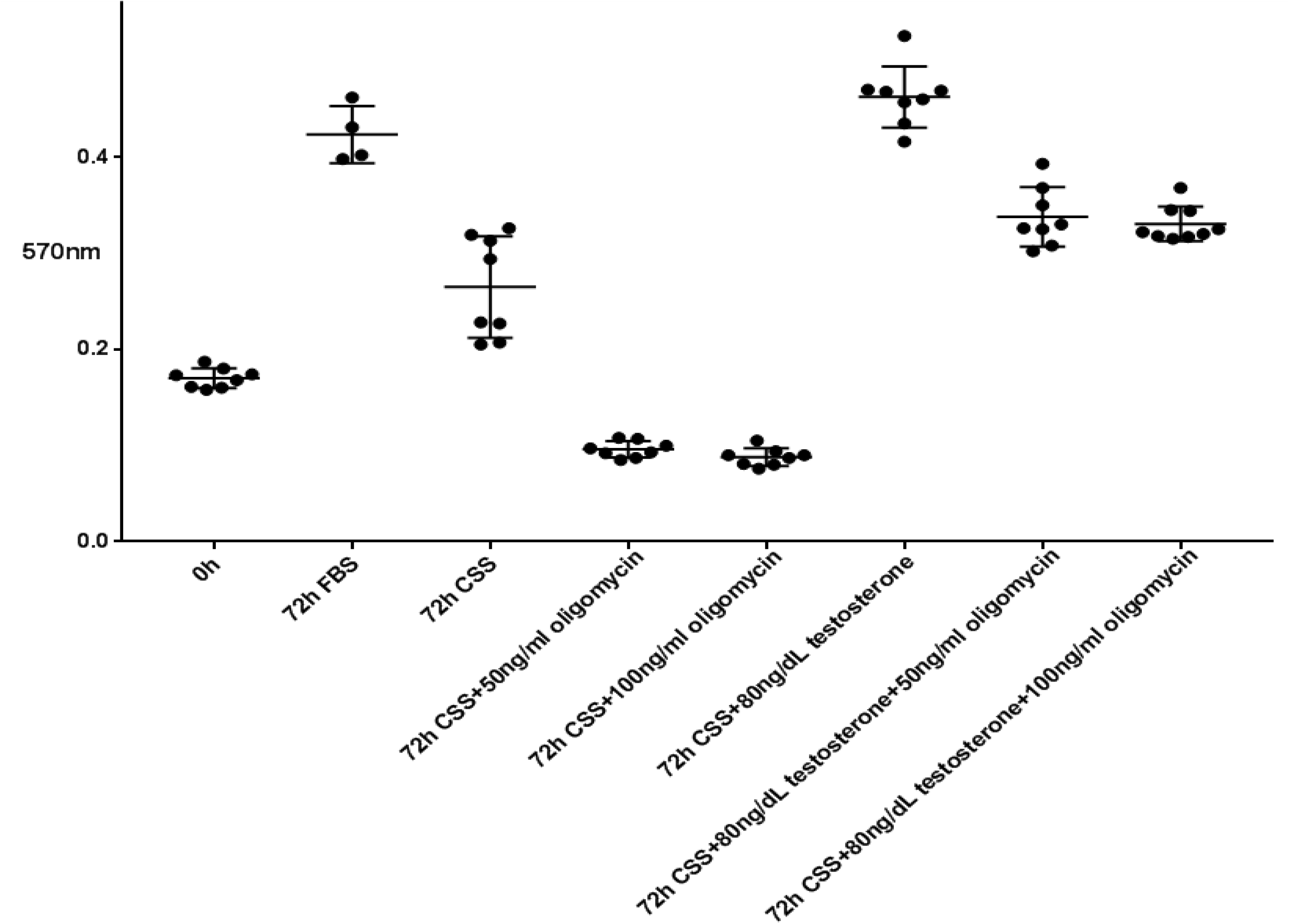

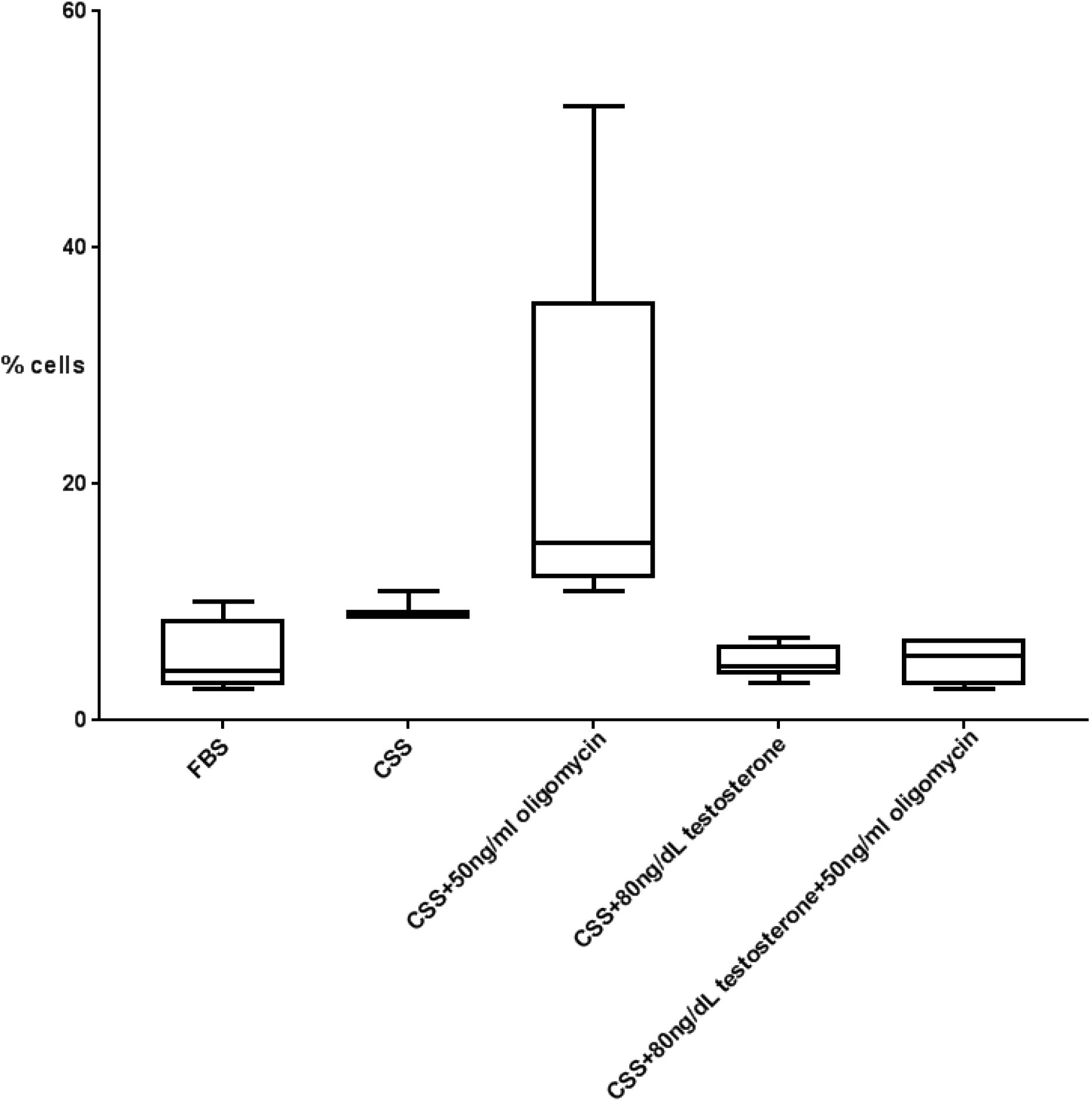

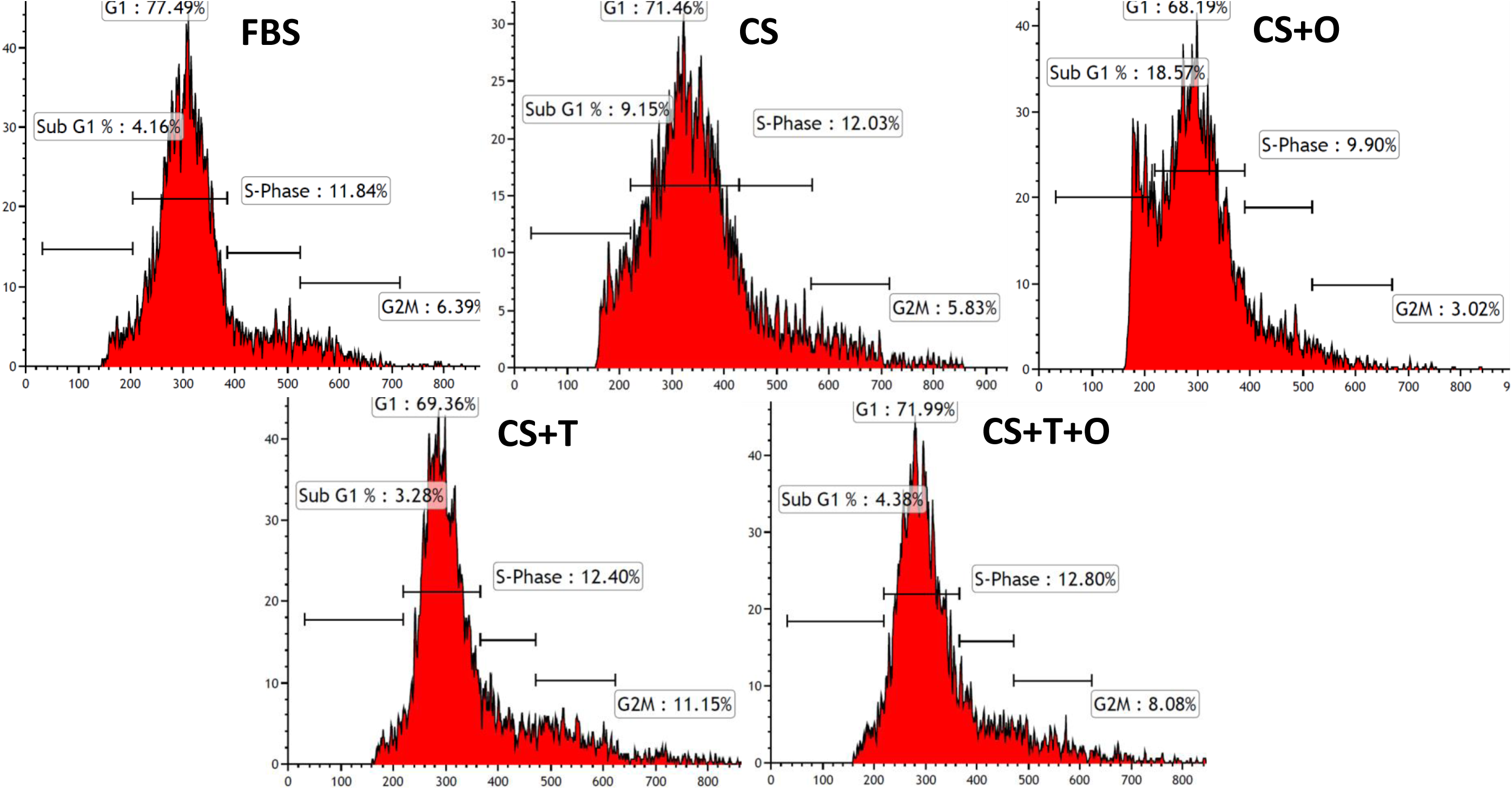

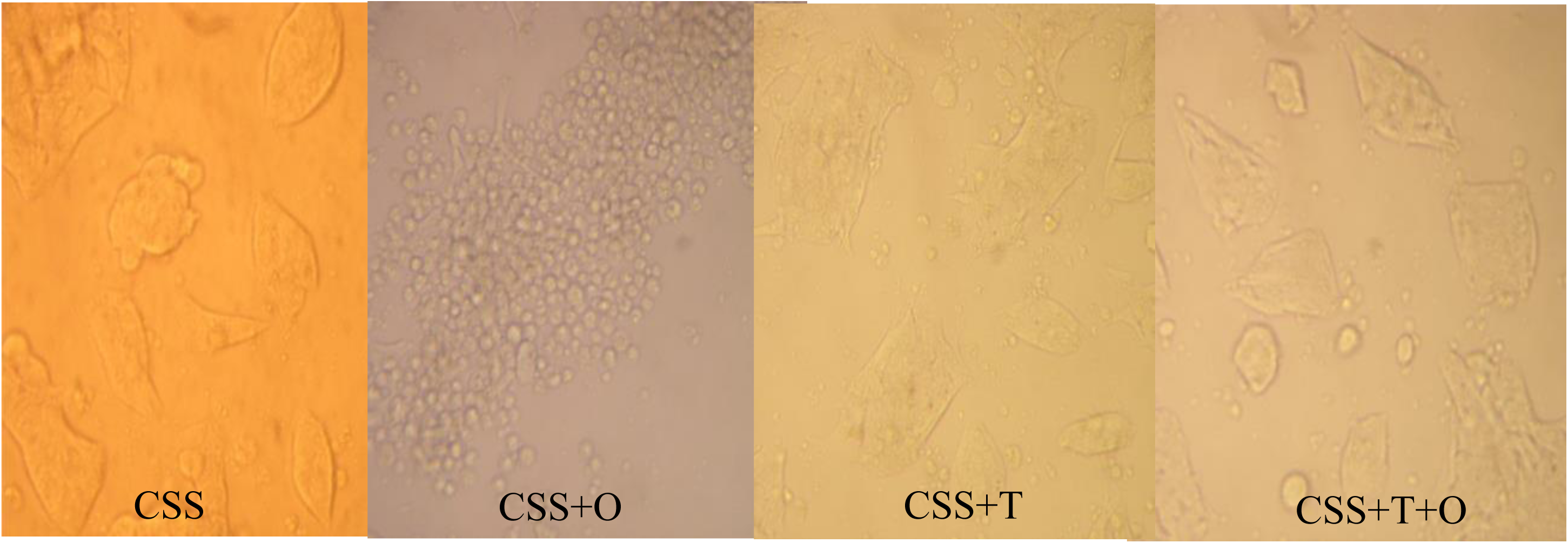

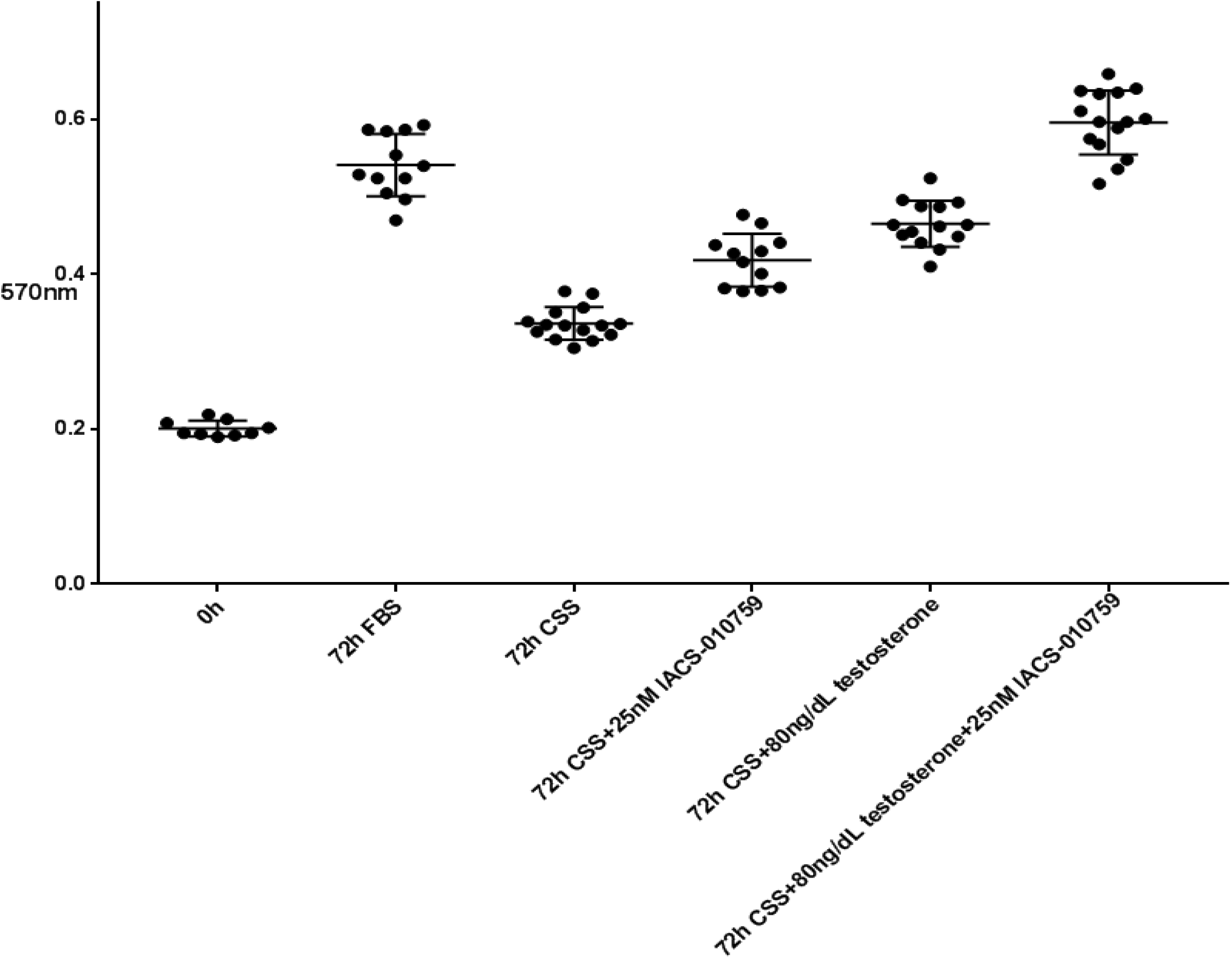

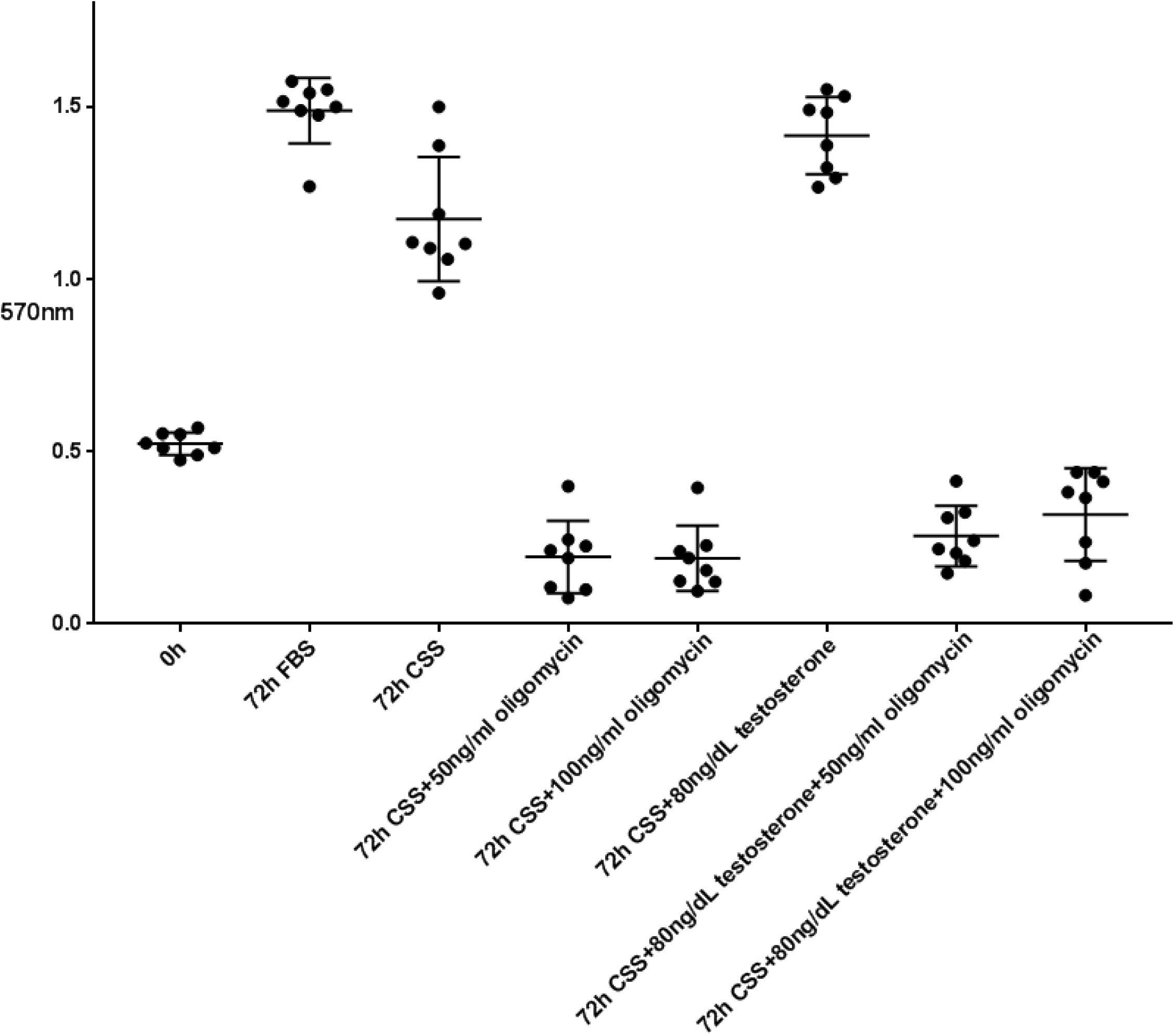

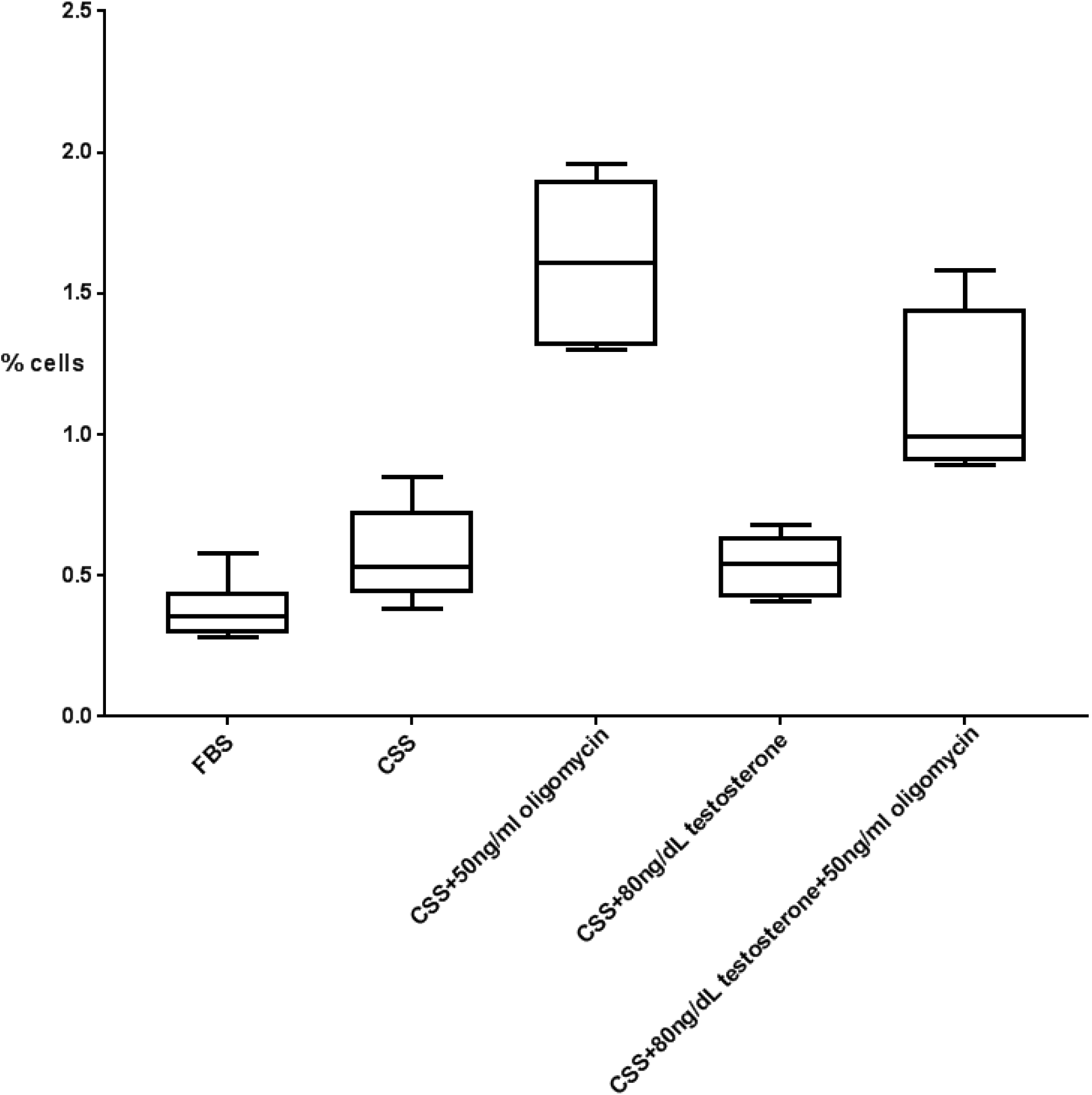

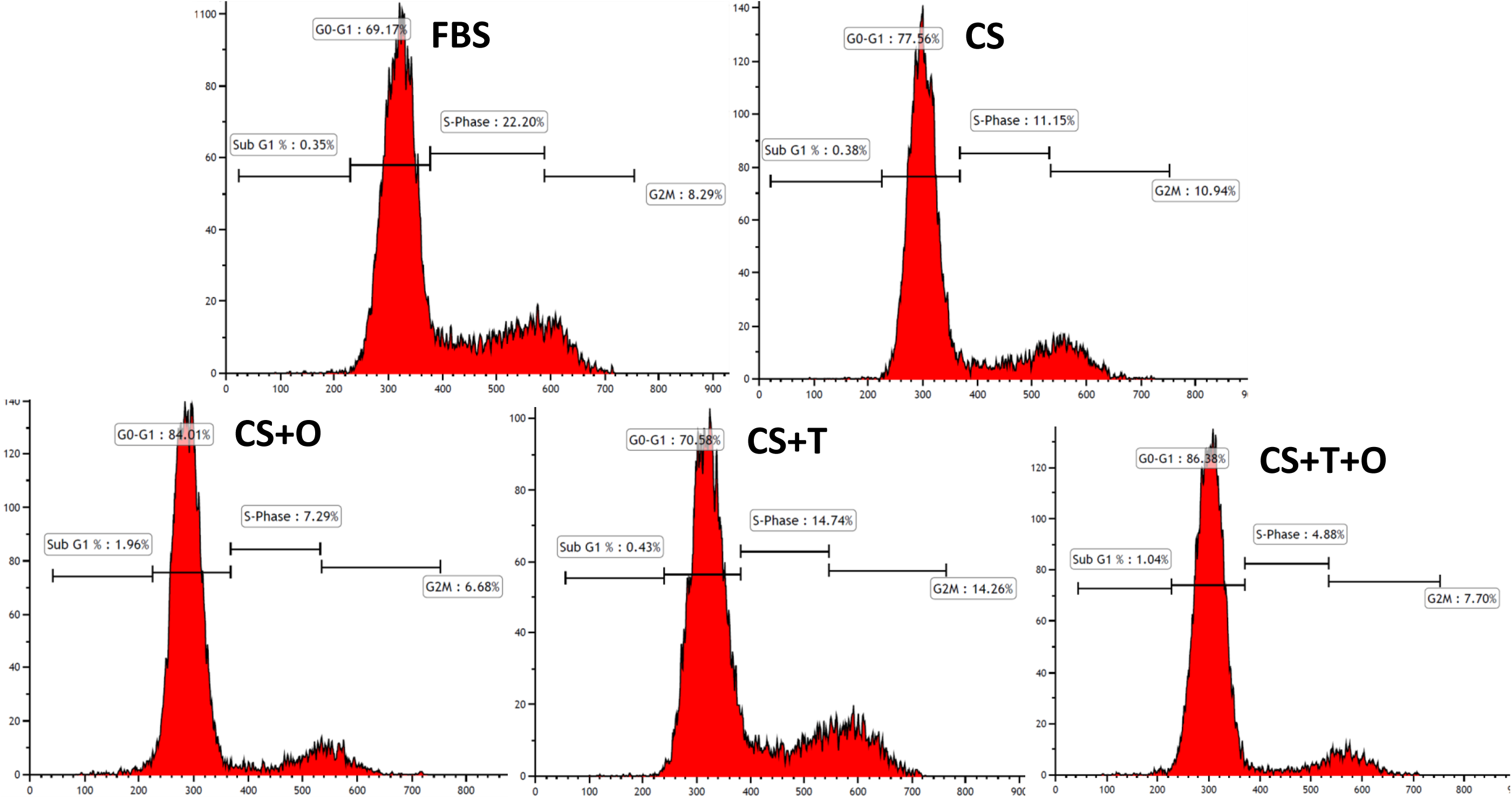

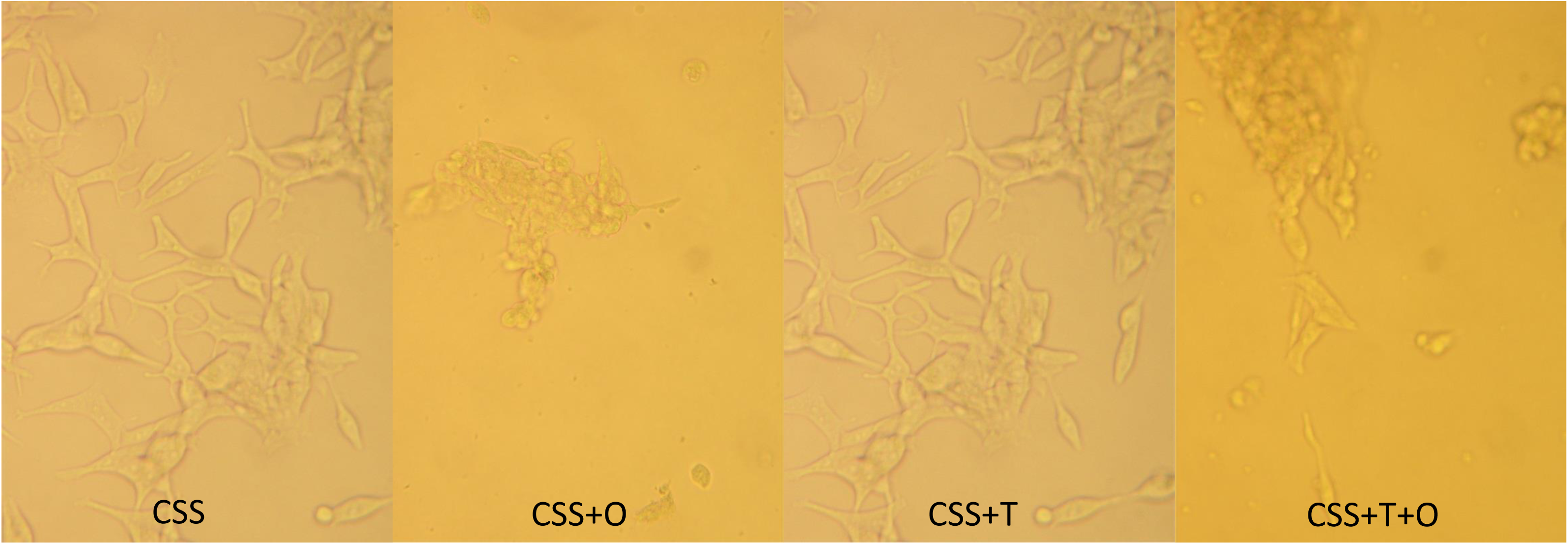

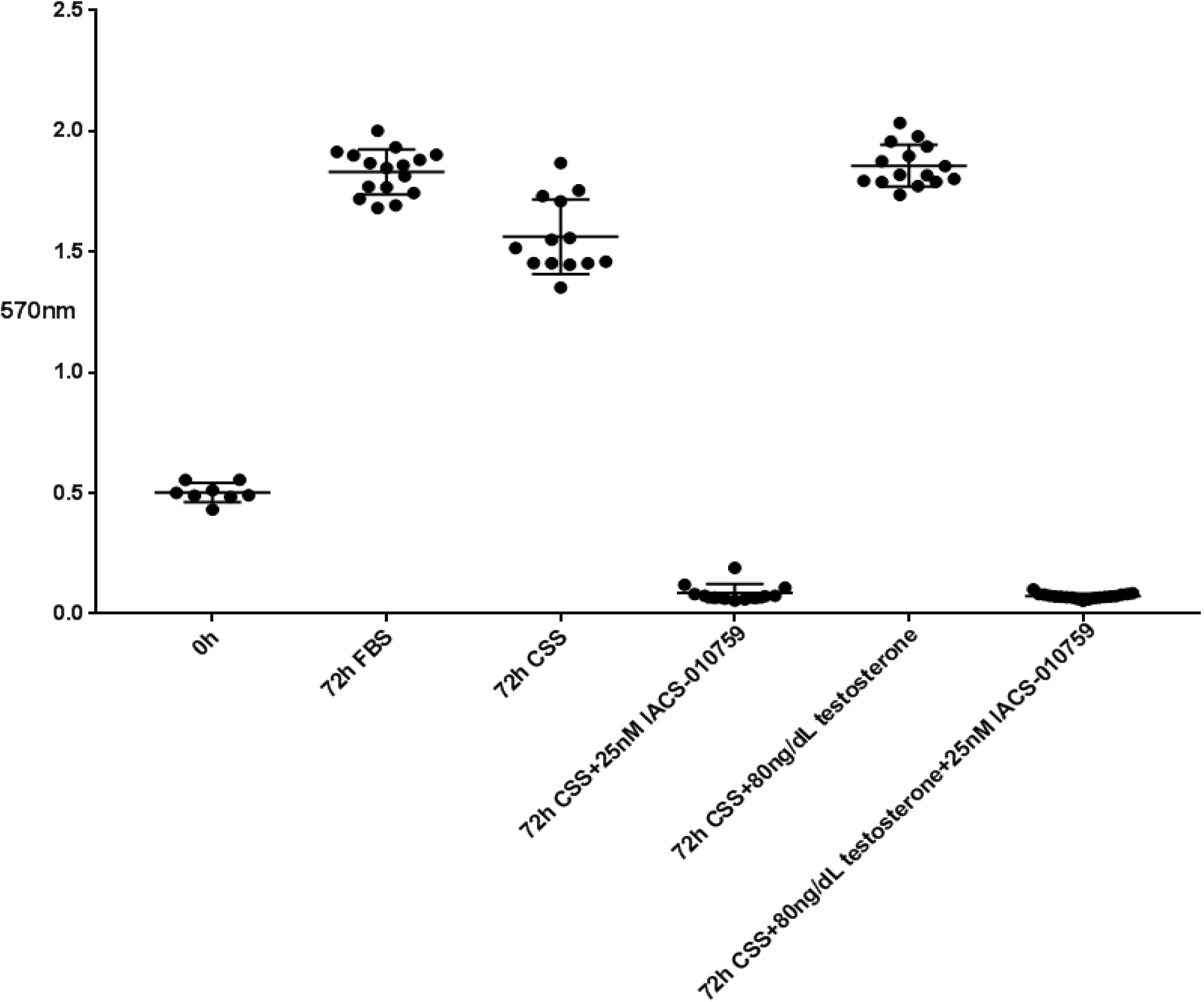

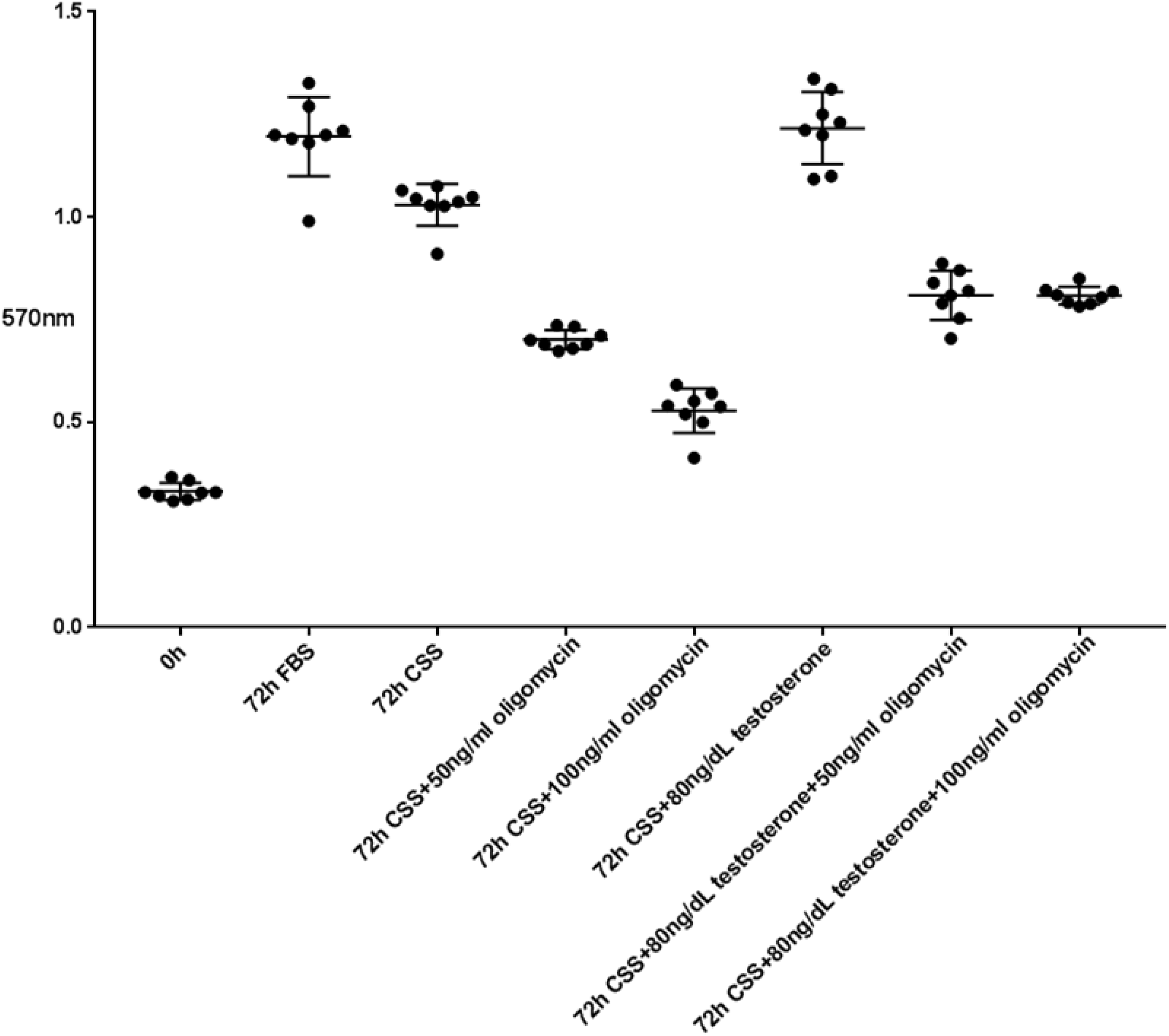

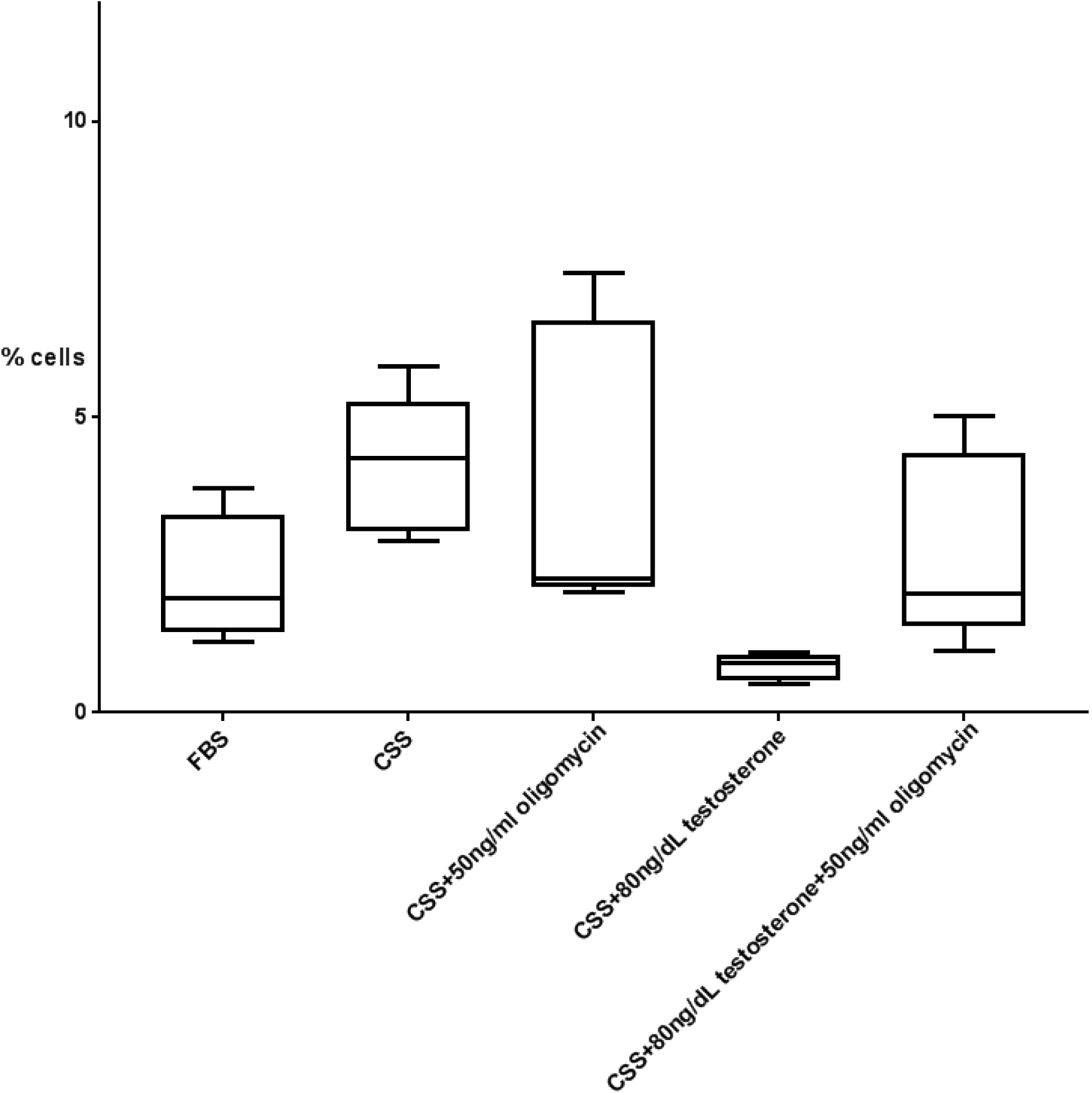

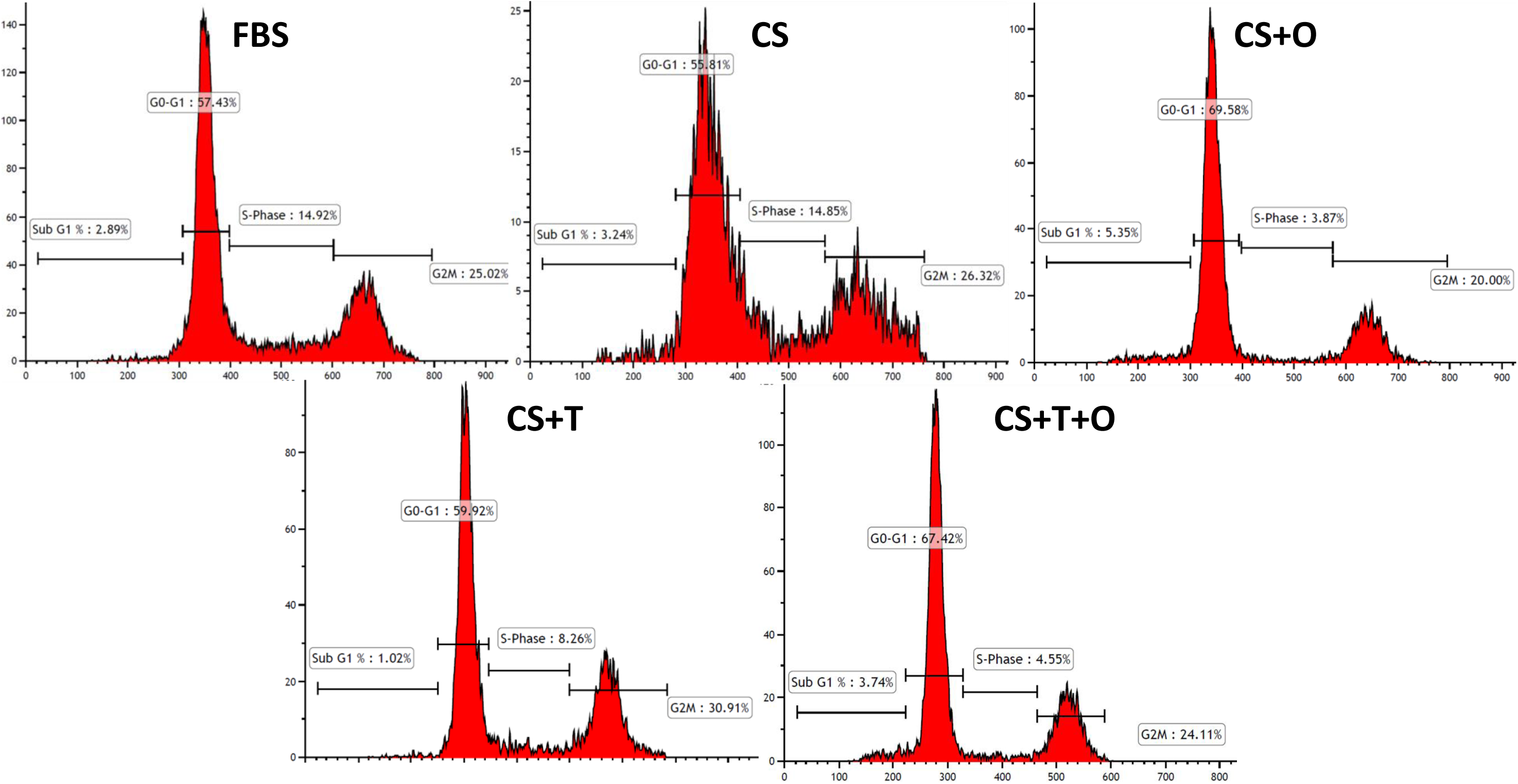

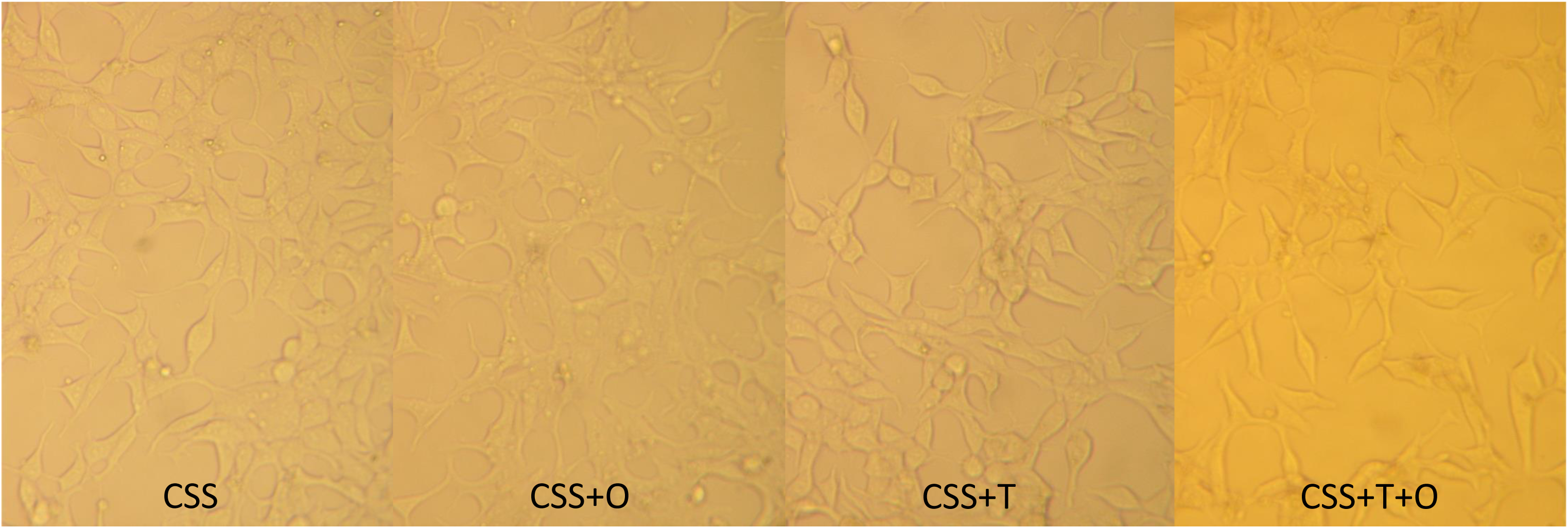

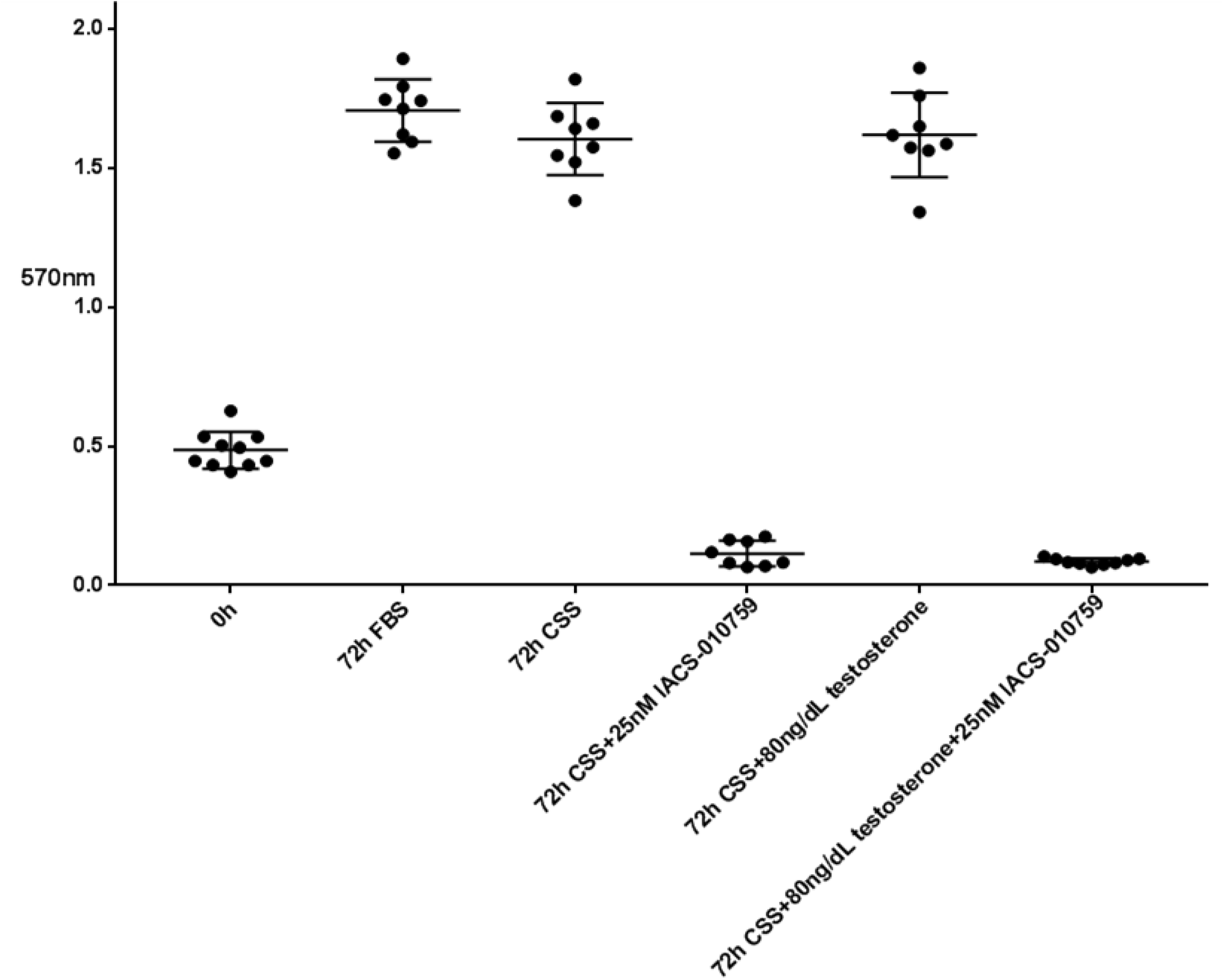

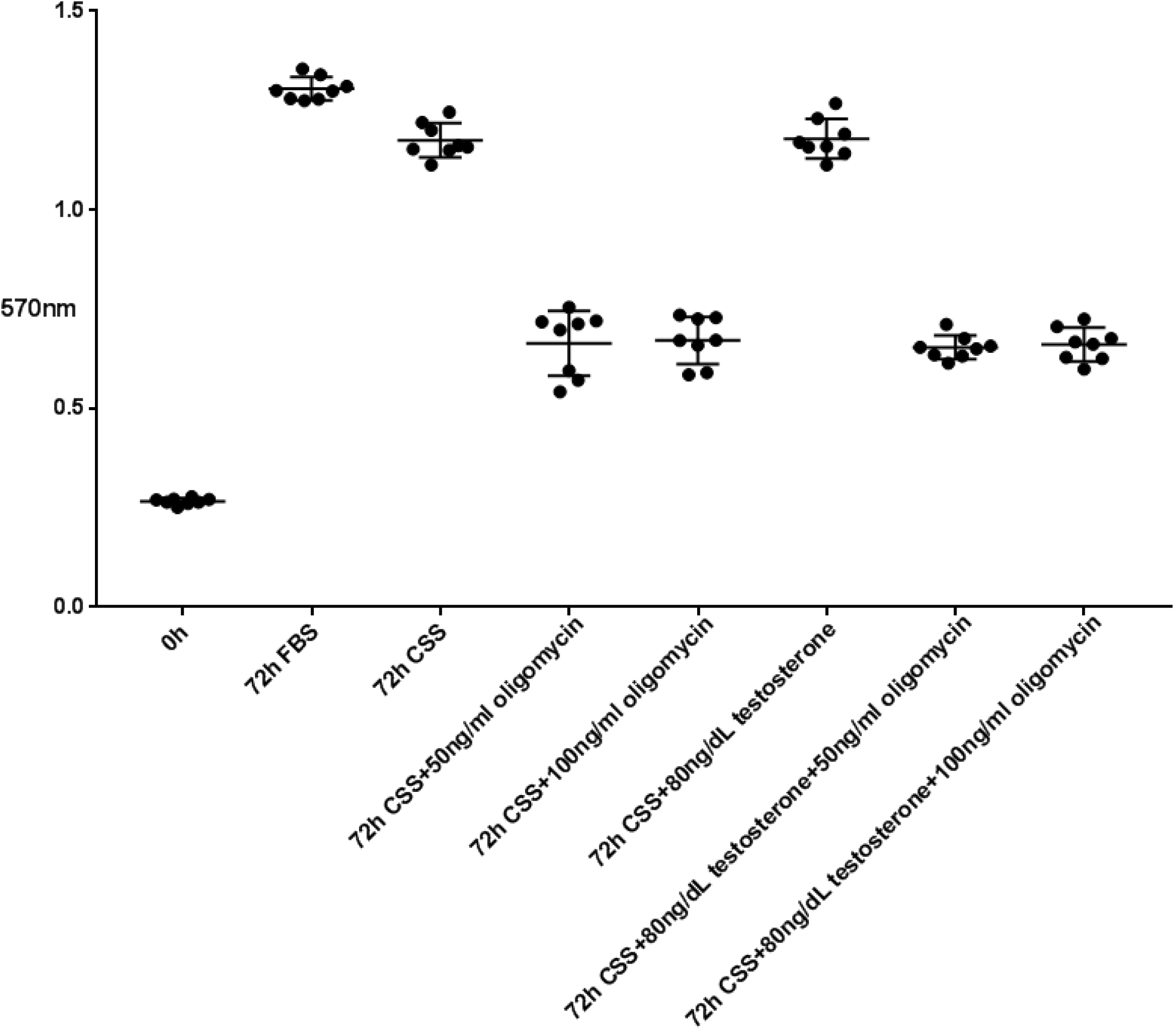

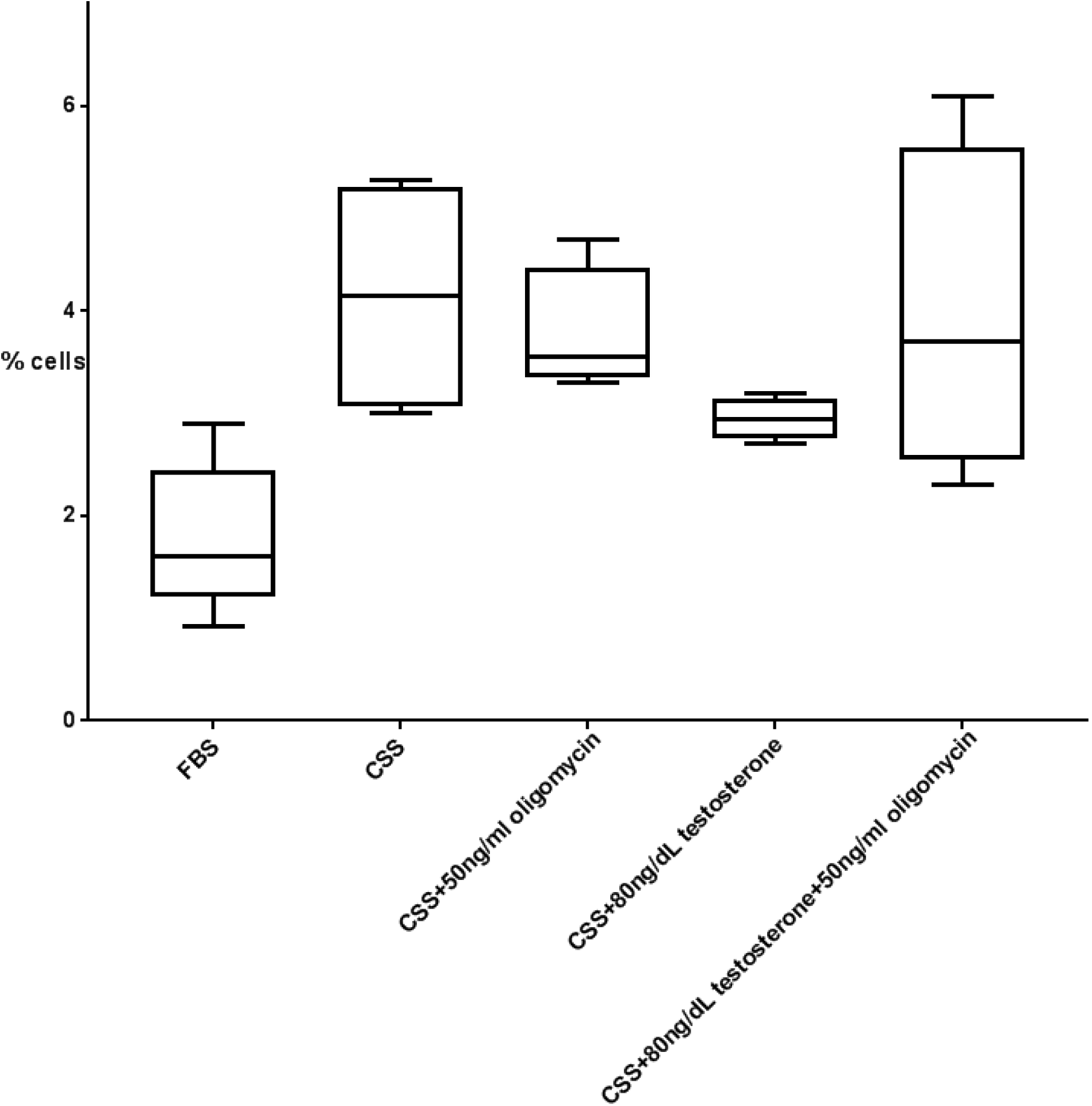

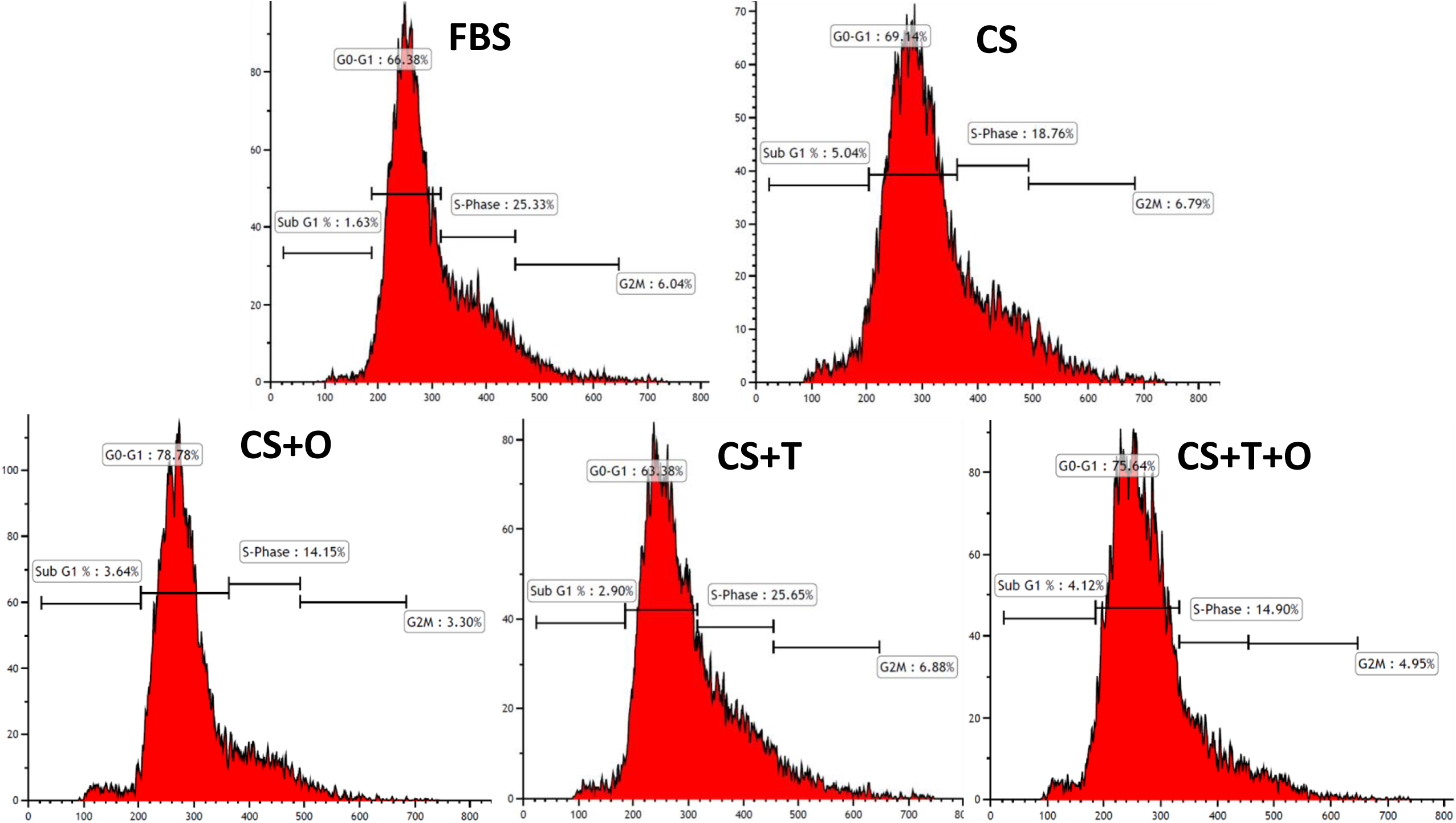

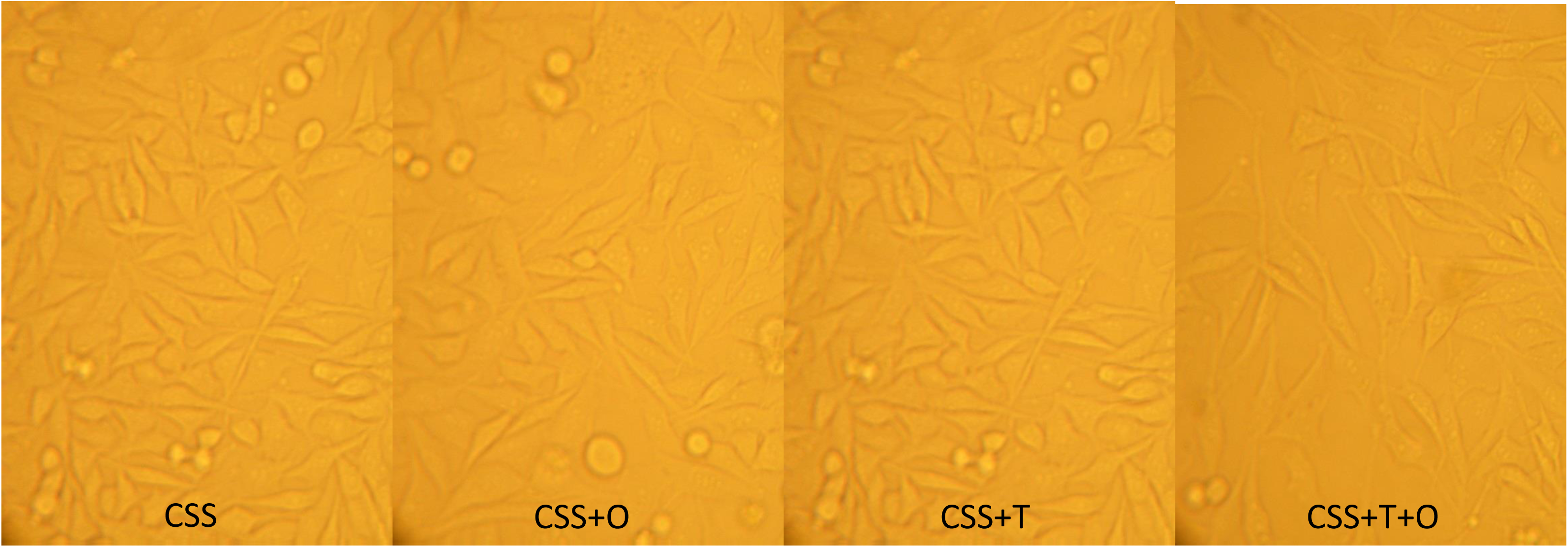

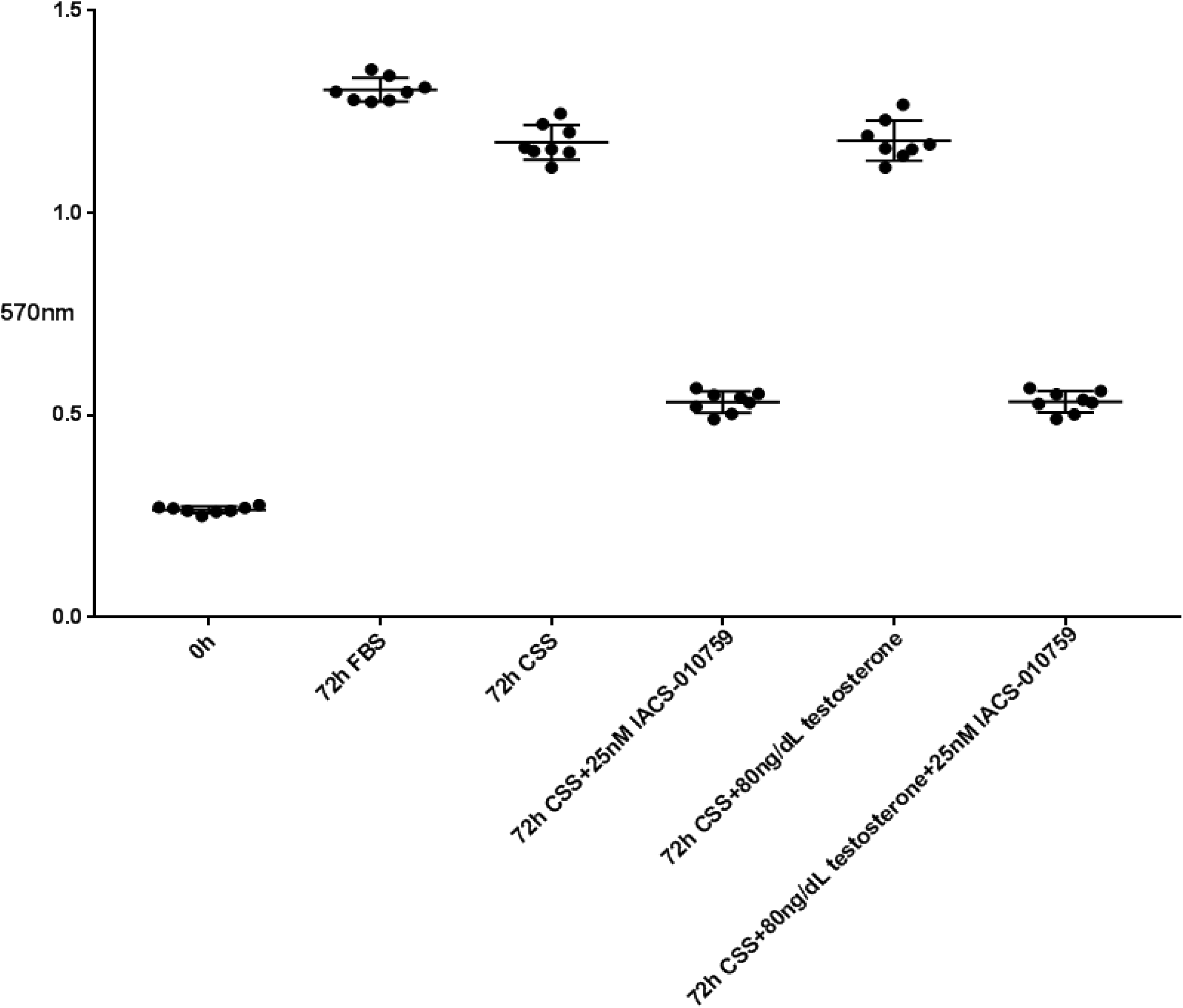

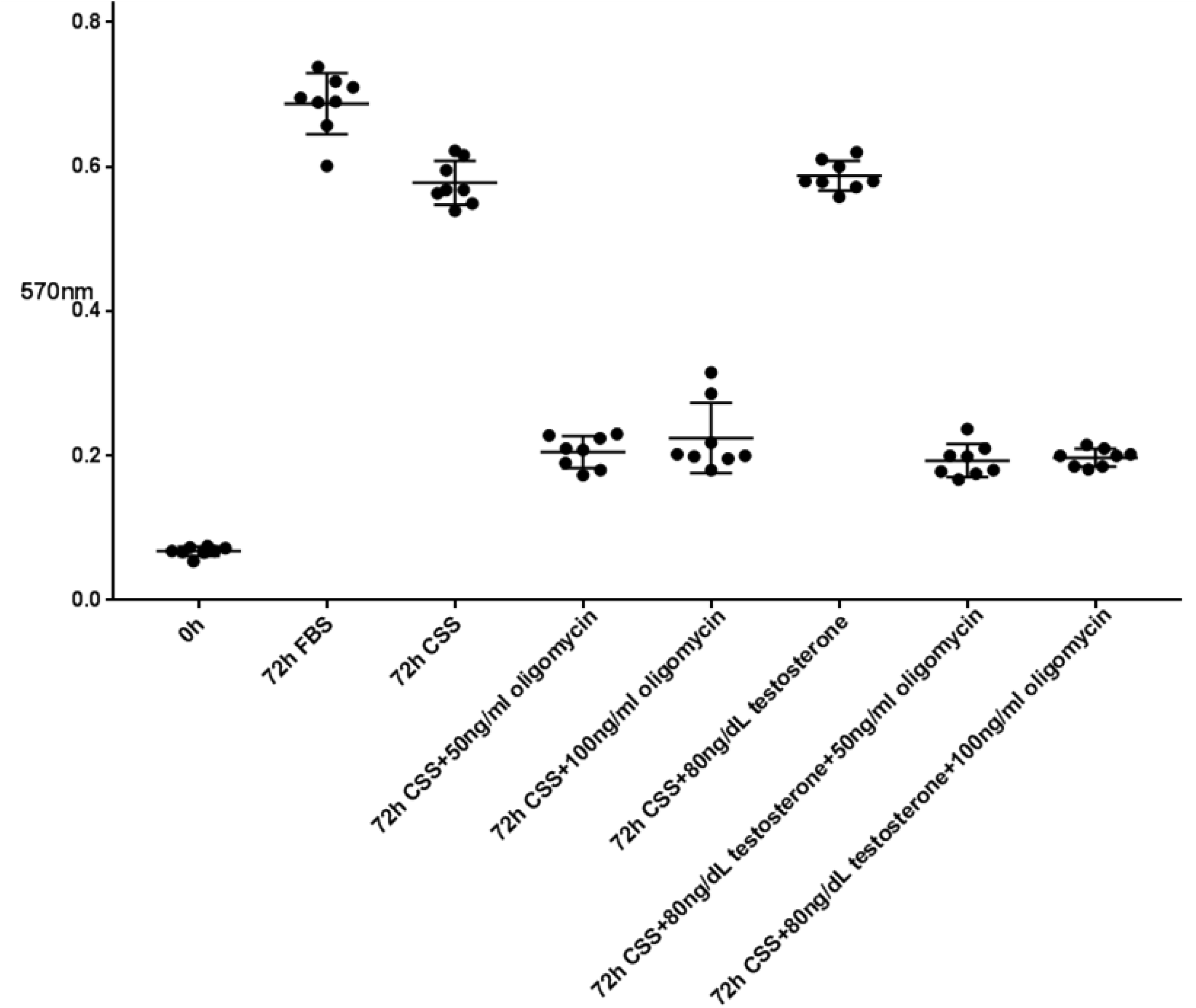

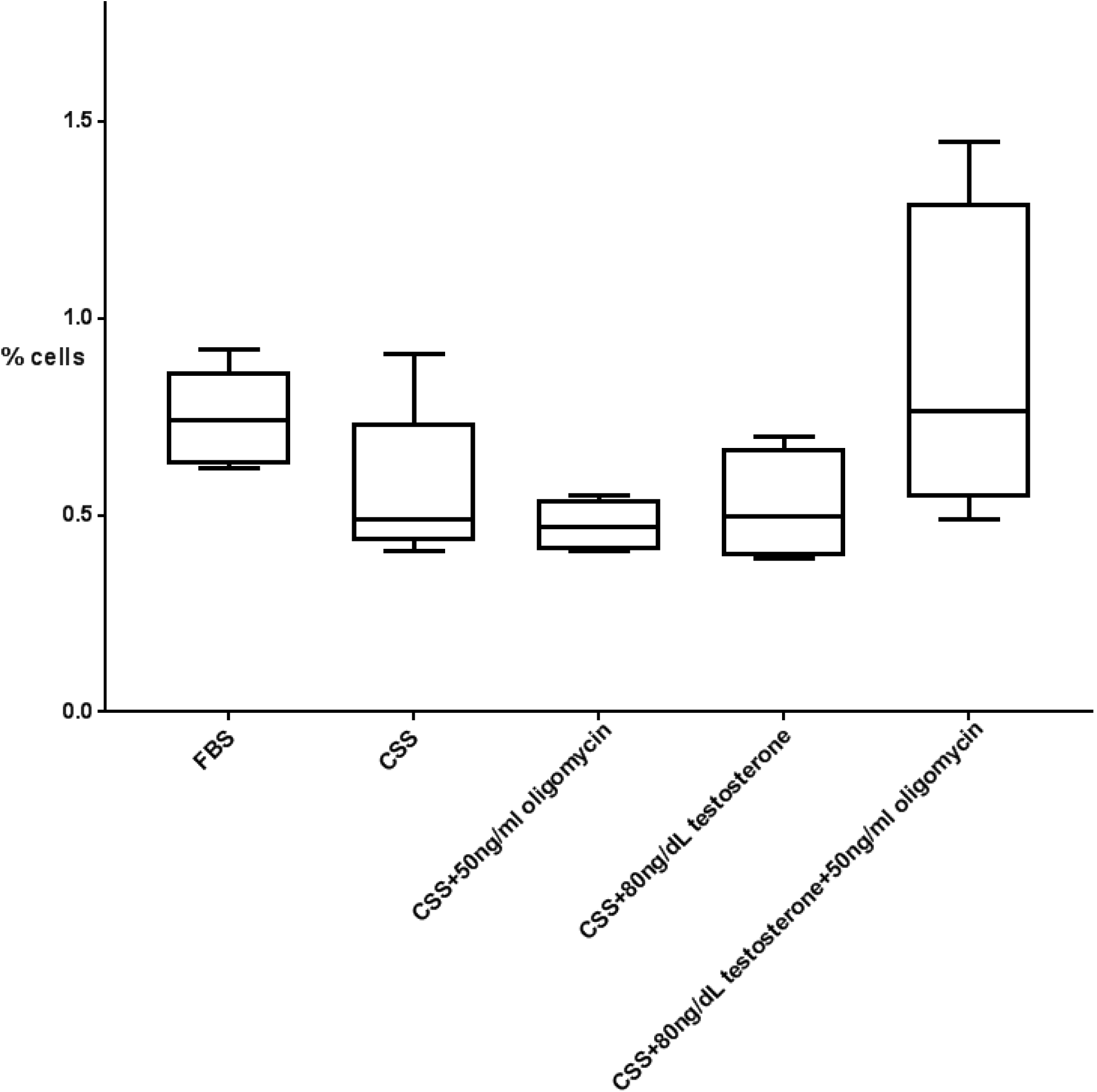

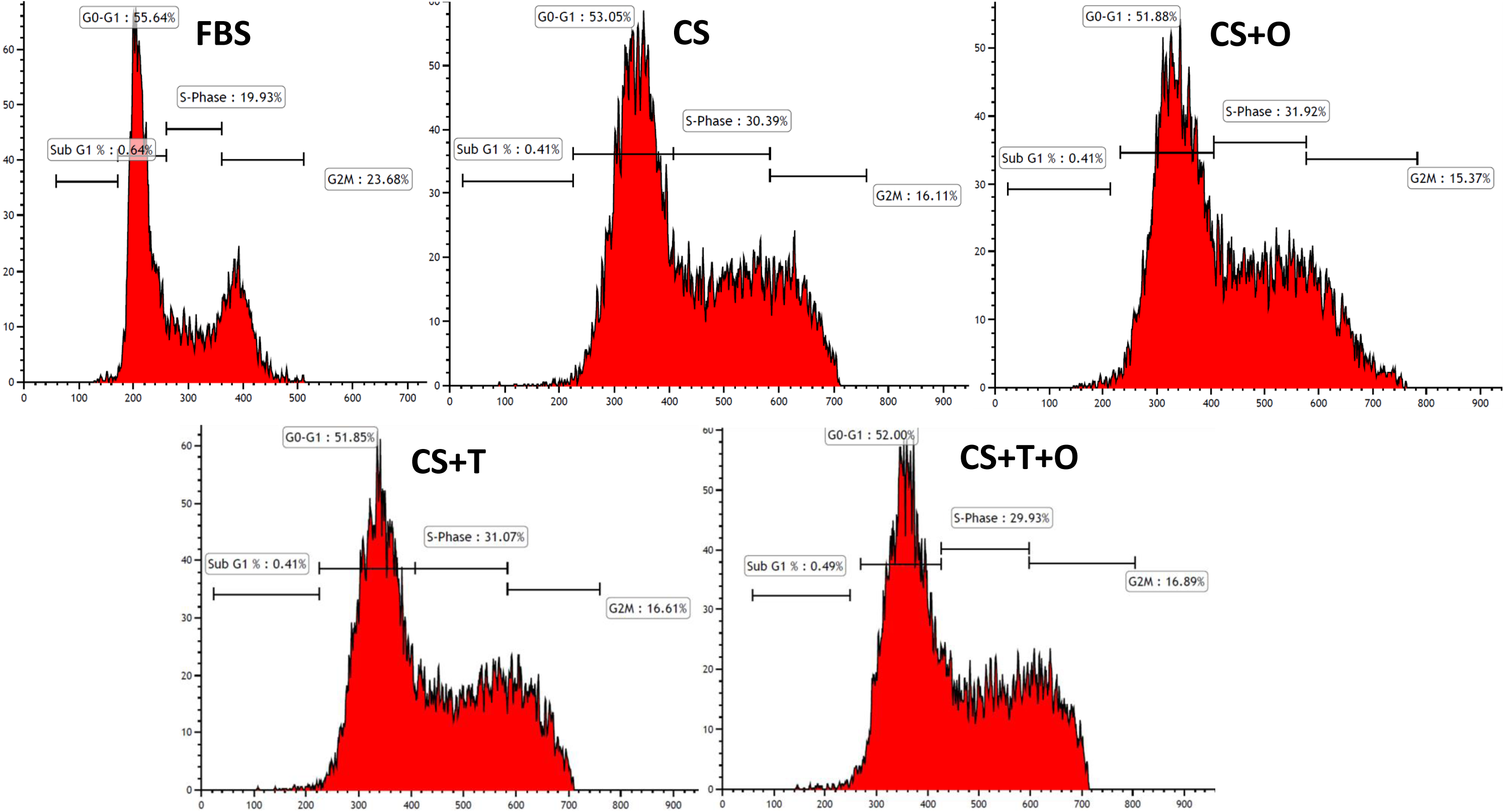

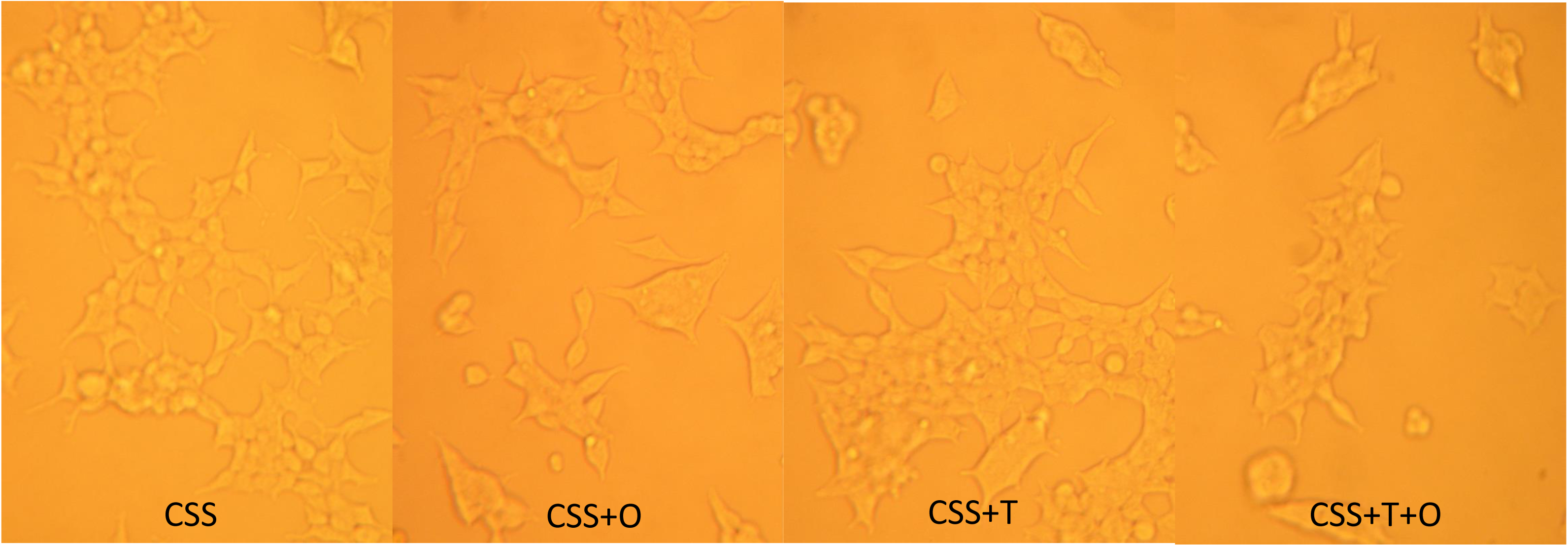

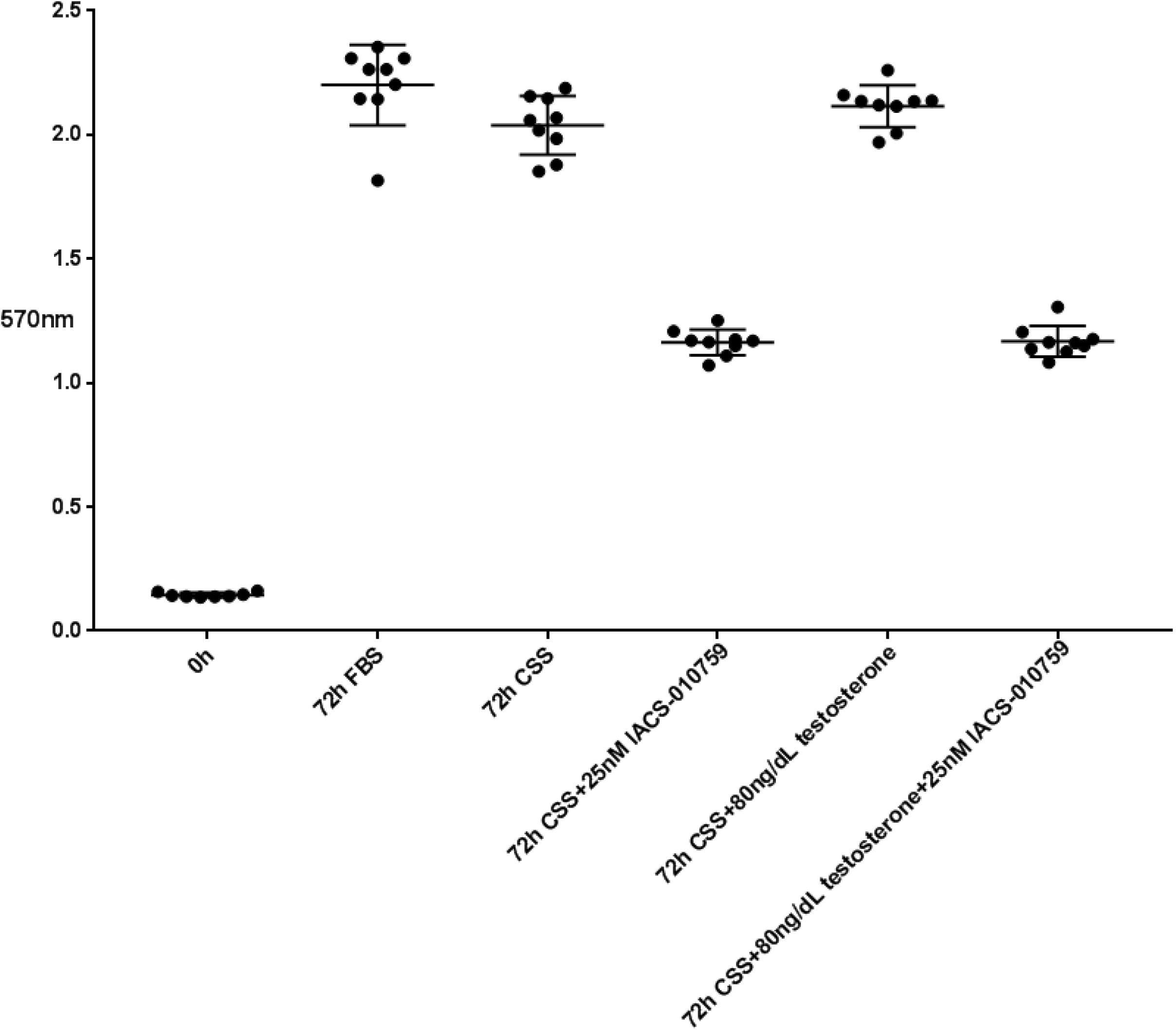

